# Social learning about rewards – how information from others helps to adapt to changing environment

**DOI:** 10.1101/2021.03.09.434563

**Authors:** M. Winiarski, J. Borowska, R. M. Wołyniak, J. Jędrzejewska-Szmek, L. Kondrakiewicz, L. Mankiewicz, M. Chaturvedi, K. Turzyński, D.K. Wójcik, A. Puścian, E. Knapska

## Abstract

Being a part of a social structure is key for survival and reproduction. Living with conspecifics boosts evolutionary fitness, by providing essential information about the environment. Nonetheless, studying neural mechanisms of social learning has not yet been established under laboratory conditions. To examine how socially passed information about the reward affects the behavior of individuals we used Eco-HAB, an automated system for tracing voluntary behavior of group-housed mice living under semi-naturalistic conditions. We show that a scent of a rewarded individual has profound effects on the conspecifics’ ability to find the reward in both familiar and novel environments. Importantly, the animals display clear and stable individual differences in social behavior. As a result, socially conveyed information has different effects on individual mice. Further, we show that disrupting neuronal plasticity in the prelimbic cortex with nanoparticles gradually releasing TIMP metallopeptidase inhibitor 1, disrupts animals’ social behavior and results in decreased ability to adapt to environmental changes. The experimental paradigm we developed can be further used to study neuronal mechanisms of social learning.

## Introduction

Social structure resulting from kinship, friendship, and hierarchy among individuals is a hallmark of human society. Being a part of this structure is key for survival and reproduction^1,2^. Similar structures, albeit less complex, are also observed in other social species, including rodents^3^. Living with conspecifics raises individual’s evolutionary fitness in many ways, among others by providing valuable information about the environment. Social learning about threats and opportunities helps individuals to adapt to the rapidly changing circumstances without the need for first-hand experience, which can be costly, especially in case of exposure to predators or energy expenditure^4,5^. Yet, studying social learning and mechanisms of socially driven spreading of information between group-housed mice is still not established under laboratory conditions.

So far processing of information provided by others has almost exclusively been studied in simplified models employing dyads of interacting rodents^6,7^. These studies showed that emotionally aroused animals transmit signals that can be detected and decoded by conspecifics^8^. They also elucidated a set of neuronal circuits involved in processing of social information, including networks within the prefrontal cortex (PFC)^9^. In contrast, how learning scales up from individuals to social networks is yet to be determined. Moreover, we know very little about social learning under more complex conditions, i.e., in animals living in groups, using larger territory and functioning within a social structure. Although difficult, such studies are of paramount importance, since mouse sociability is heavily dependent on population density, habitat size and structure^10^.

Further, most laboratory studies on social communication and learning rely on the assessment of direct interactions between individuals. However, under natural conditions, life of mice, in particular their social life, is dominated by smell. Mice commonly use odors to detect and assess food and predators, recognize individuals, and evaluate sexual and social status^11–14^. Notably, a major form of communication among mice is scent marking with urine that does not require direct contact between individuals^15,16^. Importantly, choosing ecologically relevant stimuli should increase the odds that animals master the task quickly, react in a coherent way and – crucially for brain studies – consistently use similar, well-conserved neural circuitry^17,18^.

Both rodent and human studies implicated the PFC in processing of social information^19,20^, discriminating of affective state of other individuals^21^, social hierarchy^22^, and social transmission of information about food safety^23^. Proteases have been shown to play a major role in experience-dependent plasticity both in animals^24–26^ and in humans^27,28^. In particular, TIMP-1, tissue inhibitor of metalloproteinases, has been shown to disrupt neuronal plasticity in the PFC^26^. Here we used nanoparticles gradually releasing TIMP-1 over several days^29^ to affect functionality of the prelimbic part of the PFC (PL). To study how information about the reward affects behavior of individuals we used Eco-HAB^30^, an automated system for tracing social behavior and learning of mice living in a group under semi-naturalistic conditions. Eco-HAB allows for measuring voluntary behaviors, with animals responding at their own pace, as well as for collecting data for much longer than in classical tests.

In the following studies we illustrate the profound effects of social information about reward on animals’ behavior and their ability to adapt to changes in both familiar and novel environments. We show that mice effectively learn from olfactory social cues and use that information to gain access to the reward. Importantly, the animals display clear and stable individual differences in social interactions and thus socially conveyed information has different effects on individual mice. Finally, social learning and resulting adjustments of social structure are impaired when synaptic plasticity in the PL is disrupted.

## Results

### Scent of a rewarded mouse attracts other mice and changes the pattern of their social interactions

After 48 hours of adaptation to the novel environment of the Eco-HAB system, two mice out of 12 living together (cohort 1), were separated from the group and put into individual cages for the next 24 hours. The separated mice provided a source of social olfactory information, bedding soaked with their scent, which was then presented to the rest of the group still inhabiting Eco-HAB. The two scents were presented for 12 hours, in two distant, opposite spaces within the territory (Fig 1A). To avoid mixing of the scents, olfactory stimuli were placed behind the perforated separators. Notably, the two separated mice did not come back to their original cohort in the testing phase, so there was no direct contact that might have resulted in the additional ways of communication between the conspecifics. The same cohort of mice was subjected to two rounds of experimental procedures. In the first, control round of testing, both separated mice got access to tap water while singly housed. In the second round, the same pair of mice was separated, but this time one of them got access to sweetened water (highly rewarding 10% sucrose solution), while the other could drink only tap water. We observed that during exposure to bedding from two separated animals that had access to tap water, the rest of the cohort inhabiting Eco-HAB did not prefer any of the presented scents (Fig 1B, CTRL). However, when bedding from a mouse having access to 10% sucrose solution was presented, the animals displayed a strong preference to its scent in comparison to the scent of a mouse having access to water (Fig 1B). The preference persisted for the whole testing phase, i.e., 12 hours (Fig 1C). Moreover, the presence of the scent from the mouse drinking sweetened water changed the pattern of social interactions of the tested mice, who started following each other more often (Fig 1D). Importantly, the overall locomotor activity of the mice was not changed (Fig S1), which suggests that the olfactory information affected social rather than general exploratory behavior.

**Fig 1.**
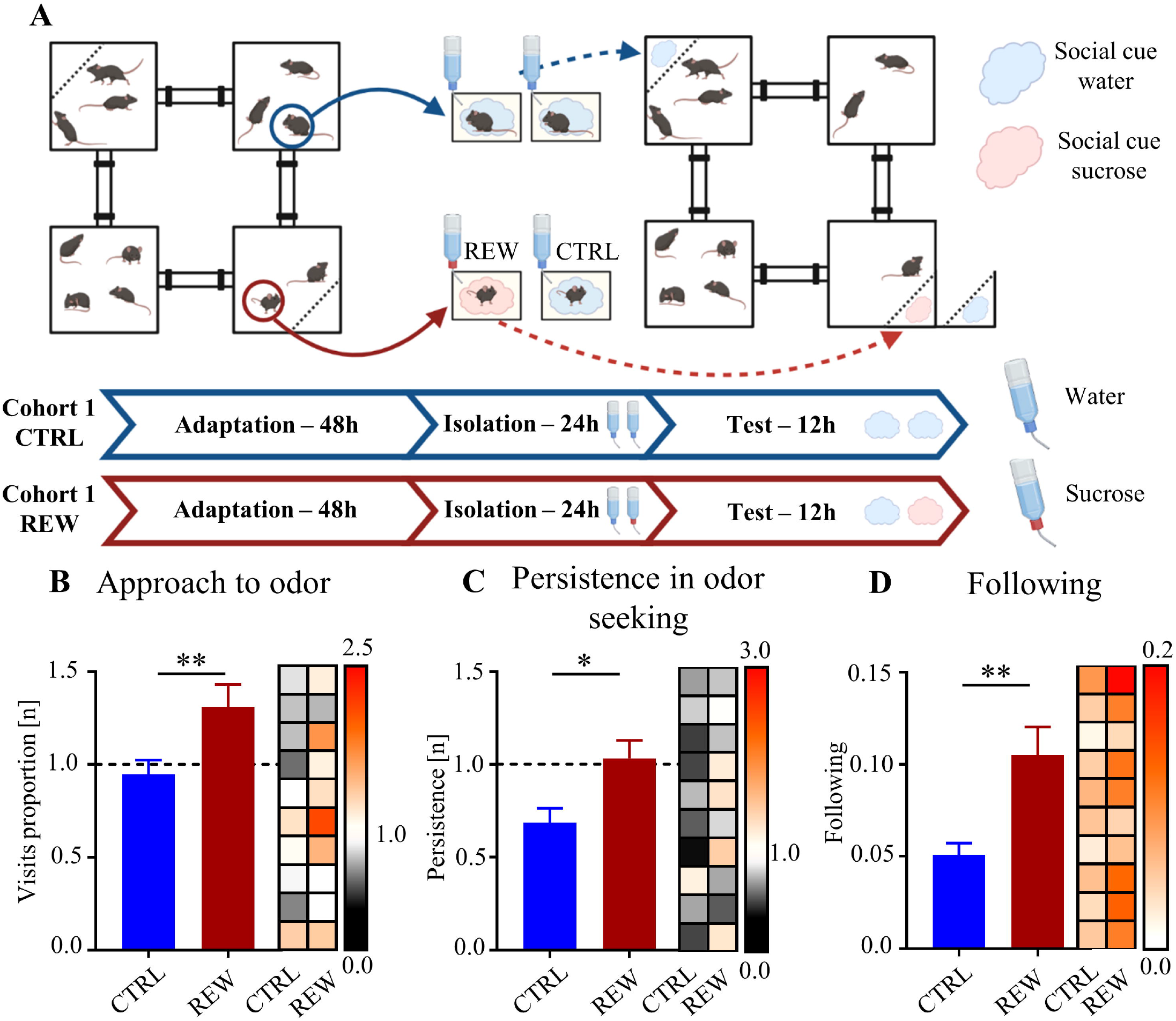
Social olfactory cues indicating reward attract mice and change the pattern of their social interactions. (A) Schematic representation of the experimental design. Arrows indicate relocation of the animals or their scents. (B) Mice preferred the compartment where the bedding from a rewarded conspecific was presented over the one from the unrewarded conspecific. Approach to social odor was calculated as a proportion of visits to the compartments containing olfactory cues (reward to neutral) divided by a respective parameter from the corresponding period 24h prior to the introduction of the social stimuli (see <u>Materials and Methods</u>). (C) The preference for the olfactory cues from a rewarded mouse persisted for the whole testing phase. Persistence in odor seeking was measured as a proportion of visits to the compartments containing olfactory cues (reward to neutral or neutral-neutral) during the second 6 hours of the testing phase divided by a proportion of visits to these compartments during the first 6 hours of the testing phase (see <u>Materials and Methods</u>). (D) Presence of the olfactory cues from a rewarded mouse increased the number of followings, in the corridors of the Eco-HAB system (see <u>Materials and Methods</u>). *p<0.05, **p<0.01, **CTRL -** control trial, **REW** - reward trial, the dotted line shows level of no change. Results of the statistical comparison to no change level are not shown to keep graph readability (B - **REW –** p<0.05, **CTRL** – n.s., C – **REW** – n.s., **CTRL** – p<0.05). Values for individual subjects are presented on the heat map (right to the bar plot), squares in each row represent data for the same mouse, columns represent trials.

### Disrupting synaptic plasticity in the prelimbic cortex impairs response to scent of a rewarded mouse

Using a variant of the above described behavioral testing in which animals were exposed to social scents indicating reward we now studied behavior of mice before (control condition) and after releasing a tissue inhibitor of metalloproteinases TIMP-1 into their prelimbic cortex (PL, Fig 2A). Specifically, at first, we tested a new cohort of naïve mice (cohort 2) and replicated the previous finding, that is a preference for the social odor indicating reward (Fig 2B). Then we bilaterally injected animals with nanoparticles loaded with TIMP-1 to the PL, which significantly changed the observed behavioral pattern. Brain-manipulated mice showed no persistent preference for the scent of a rewarded mouse (Fig 2C), and followed each other significantly less (Fig 2D). Importantly, injection of the TIMP-1 nanoparticles did not change the overall activity, indicating social specificity of the observed impairment (Fig S2).

**Fig 2.**
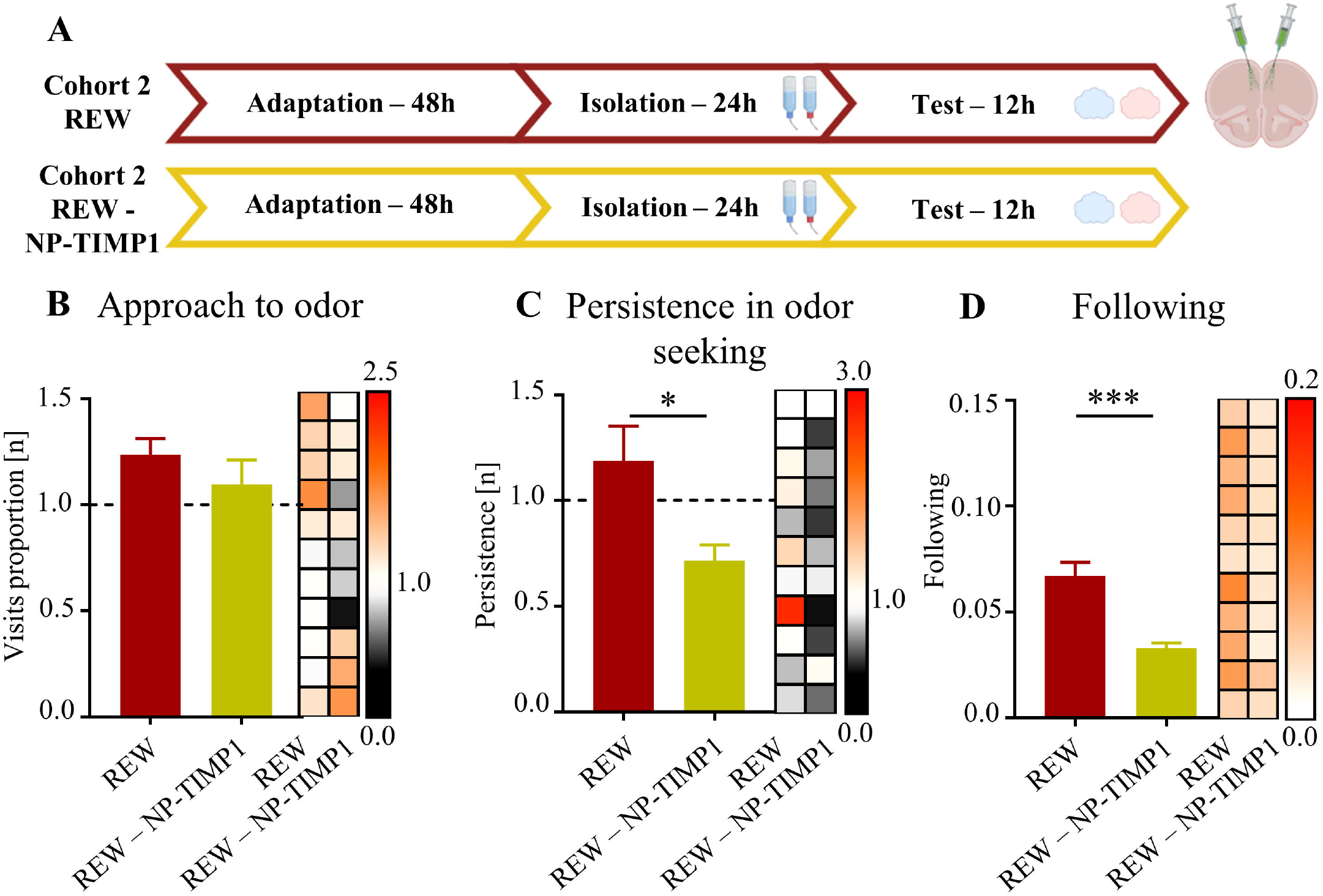
Disrupting synaptic plasticity in the prelimbic cortex impairs response to olfactory cues from a rewarded mouse. (A) Timeline of the experimental protocol. First, we tested response of mice to the olfactory cues from a rewarded conspecific (**REW**), then the mice were stereotaxically injected with nanoparticles gradually releasing TIMP-1 (NP-TIMP1) into the prelimbic cortex (PL) and, after 5 days of recovery, retested in the Eco-HAB (**REW – NP-TIMP1**). (B) TIMP-1 injection to the PL did not change the preference for the olfactory cues from a rewarded mouse when compared to the trial before surgery. However, it decreased persistence in its exploration (C) and followings within the tested cohort (D). All the measures were calculated as for the data presented in Fig. 1. *p<0.05,*** p<0.001, the dotted line shows the level of no change. Results of the statistical comparison to no change level are not shown to keep graph readability (B - **REW –** p<0.05, **REW-NP-TIMP1** – n.s., C-**REW –** n.s., **REW-NP-TIMP1** – n.s). Values for individual subjects are presented on the heat map (right to the bar plot), squares in each row represent data for the same mouse.

### Social olfactory information helps to find the reward in a novel environment, which requires an intact prelimbic cortex

To investigate whether olfactory information from a rewarded mouse helps to find reward in a novel, previously unknown environment we tested mice (cohorts 3, 4 and 5) transferred to a different Eco-HAB system, previously inhabited by two familiar mice, who had left social olfactory cues throughout the territory. Said social cues had been left by the animals that had access to either tap water in two opposite Eco-HAB compartments (control condition - cohort 3) or to sweetened water (10% sucrose solution) in one of the compartments and to tap water in the other compartment (reward condition – cohorts 4 and 5). Importantly, the bottles were replaced with the new ones containing tap water before the tested cohort of mice, moving in from another Eco-HAB, was introduced to the testing environment (Fig 3A). In addition to the previously described behavioral measures we assessed the time spent on drinking water from both bottles and observed a preference for the previously rewarded site (cohort 4). Interestingly, this effect was also observed for the control group (cohort 3), in which 2 mice who had lived in the system before showed slight preference for one of the bottles, even though both contained tap water (Fig 3B). However, this effect was transient. Otherwise, the preference for the previously rewarded site persisted for the whole testing phase (Fig 3C). Further, similarly as in the previous experiments, mice followed each other more frequently (Fig 3D), when the social information about the reward was present in the environment. On the other hand, mice treated with TIMP-1 in the PL (Fig 3B-D, cohort 5) show a decreased response to information about the reward. As before, there was no difference in general activity between the control (cohort 3) and reward vehicle group (cohort 4), while TIMP-1 treated group (cohort 5) showed a slightly decreased locomotor activity (Fig S3).

**Fig 3.**
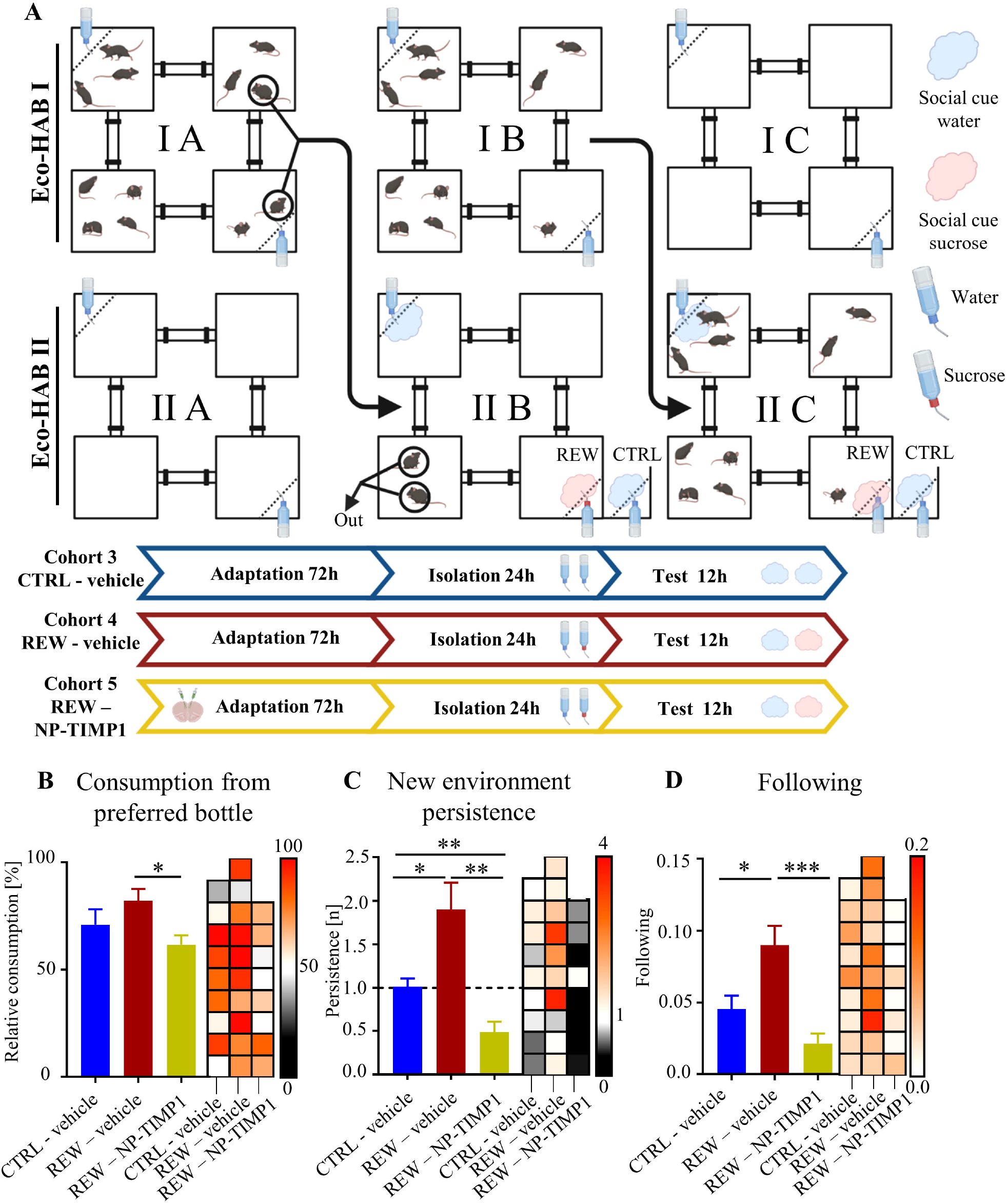
Social olfactory information helps to find the reward in a novel environment but only in mice with an intact prelimbic cortex. (A) Schematic representation of the experimental design. Two cages were used, I and II, with three phases: adaptation (A), isolation (B), and test (C). During adaptation, in the corners of two cages (**I, II**), additional bottles containing water were placed. To measure the time of drinking of individual mice an RFID antenna was mounted in front of each bottle nipple. After 72 hours of adaptation (I-A), two mice from cage I were moved to the other, clean and intact Eco-HAB II **(II-B)** leaving the rest behind **(I-B)**. One of the bottles in the novel Eco-HAB II environment contained 10% sucrose solution, the other one tap water (in the control group both bottles contained tap water). After 24h the two mice were moved out of the experiment, the bottles were cleaned and refilled with tap water, and the rest of the group was moved into this Eco-HAB II environment from cage I (**II-C)**. (**B)** Newcomer mice preferred drinking from the bottle preferred by the mice previously living in the Eco-HAB. Injection of TIMP1 decreased that preference (see <u>Materials and Methods</u>). (**C)** The preference for the bottle increased over time when reward-related social information was present. In contrast, the TIMP1 injected mice decreased the preference with time. Persistence was calculated as in the previous experiments but time spent on drinking from the bottles was used instead of the visits to the compartments and normalized to time spent in boxes with bottles (see <u>Materials and Methods</u>). (**D)** As previously, followings were increased in the REW - vehicle group, while mice treated with TIMP-1 in the PL showed decrease in following each other. *p<0.05, **p<0.01, ***p<0.001. Results of the statistical comparison to no change level are not shown to keep graph readability (C – **CTRL-vehicle** – n.s, **REW-vehicle –** p<0.05, **REW-NP-TIMP1** - p>0.05). Values for individual subjects are presented on the heat map (right to the bar plot), squares in each row represent data for the same mouse.

### Mice form stable social networks reflecting individual differences in responding to social information about reward

Getting olfactory cues through sniffing other individuals is an important source of information in mice^31^. Thus, to track social interactions that can provide such information we focused on the patterns of following between the animals cohabitating the Eco-HAB, specifically, the manner in which they trail one another through the corridors connecting different parts of the territory. We chose to measure following, since it is a type of social interaction by definition voluntarily initiated by the mouse which chooses to trail another and thus get direct access to the olfactory cues of a trailed individual. As shown by the already presented data, on average, the number of followings within each tested cohort increased when social olfactory information about reward appeared in the environment. To look into the individual differences in their intensity we visualized social network structure with a node - edge graph. In the graph, the nodes represent individual mice, and the thickness of the edge connecting two nodes represents the number of followings a given follower performs after a given leader (Fig 4A, B). We observed that mice gradually develop stable and complex social network, with differences in the level of followings between individuals (Fig 4C-H, cohort 1). The individual differences in sociability were reflected especially in the variable increase in the followings when the reward-related social cues were present in the environment (Fig 4F-H). Moreover, longitudinal observation of the social network (cohort 6) shows that it is formed very early in the experiment and remains stable throughout (Fig S4A, B). However, it is clearly disturbed when the PL plasticity is impaired with TIMP-1 (Fig S4A, C), which suggests that constant updating of information in these neuronal circuits is needed to maintain the group structure. Notably, social network distribution in mice tested in Eco-HAB shows similarities with networks observed in human research^32^.

**Fig 4.**
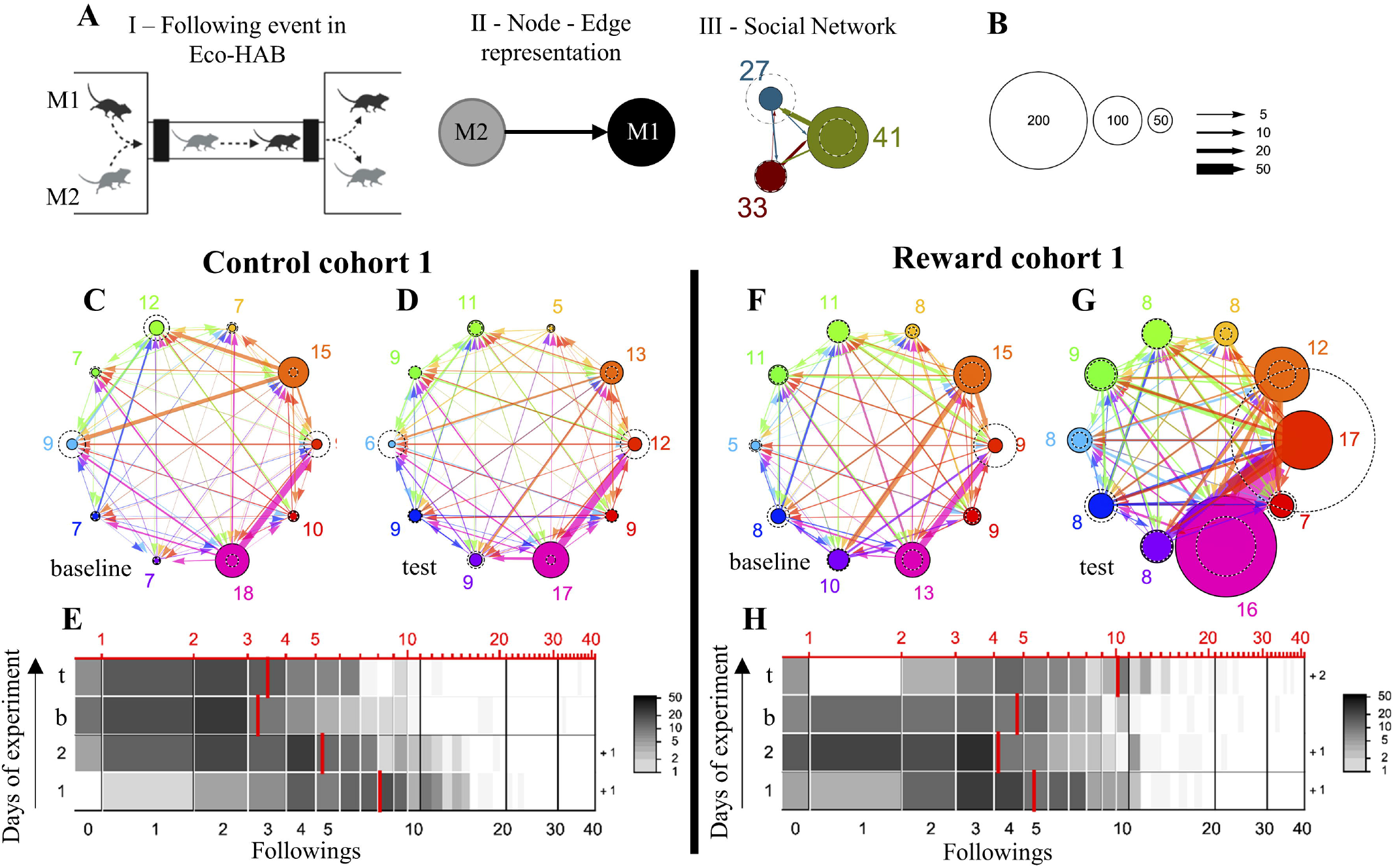
Mice form stable social networks based on individual differences in following each other. **(A)** Dynamic social interactions can be defined based on patterns of leadings/followings, i.e. (I) instances of a follower mouse (M2, gray) trailing a leader mouse (M1, black), that is passing through the corridor very shortly after it, so that the animals remain in close contact. (II) Those events form group’s social network represented as a weighted, directed graph with nodes corresponding to individual mice and edges to interactions between them. (III) Different colors represent the followings of each individual mouse. The numbers next to each node give the PageRank centrality (see <u>Materials and Methods</u>) of that node calculated for the weighted, directed graph represented by the adjacency matrix of followings; the PageRank centrality is expressed as a percentage (rounded to the closest integer), characterizing each mouse in relation to a whole social group summing up to 100%. **(B)** The radius of the colored circle at a given node is proportional to the number of followings performed by the corresponding mouse, while the radius of the dashed circle is proportional to the number of leadings performed by that mouse. The arrows are directed from a follower to a leader; the thickness of an arrow is proportional to the number of followings a given follower performed after a given leader. **(C-H)** Social network of cohort 1. observed during baseline period and after the presentation of social cues carrying information about either the neutral stimulus (C, D; scent of a naïve, unrewarded animal) or the reward (F, G; scent of a rewarded animal). Graphs show increase in following resulting from the presentation of the social cue indicating reward (F, G) and its absence in the control condition (C, D). Horizontal segments (E, H) correspond to the distribution of the number of following events in all pairs of mice within cohort 1. in the subsequent dark phases of the experiment (on days 1-4, where day 3. represents the baseline period (b) and day 4. the period of test when stimuli carrying social information were presented in the environment (t)). Subsequent days of the experiment correspond to respective rows, from bottom to top. Rectangles running from left to right reflect the number of followings ranging from 0 to 40. The intensity of the shading represents the number of occurrences of a given number of followings in all pairs of mice. In cases when there were pars of mice with over 40 followings, the number of such pairs is shown with a plus sign to the right of a respective row. Additionally, the mean number of followings for all pairs of mice during a given experimental phase, calculated for this distribution, is marked as a red line at the position corresponding to the red scale at the upper edge of the plot. As shown in (E) the intensity of following among mice is high in the beginning of experiment when animals are introduced to the Eco-HAB environment for the first time and thus explore it intensively. Notably, (H) this effect is absent when the same group of animals is reintroduced to the already familiar experimental environment for second time.

Social structures in rodents are most commonly related to social hierarchy^33–35^. Thus, to investigate the relationship of the social network based on intensity of following conspecifics with dominance hierarchy, we compared the number of followings performed by the individual mice with their performance in the U-tube dominance test (cohort 7). We show that in the well-stabilized social network there is a positive correlation between the number of followings and U-tube winning score (Fig S5).

## Discussion

We show that information about reward is coded in social olfactory cues left by a rewarded mouse. Mice are able to distinguish between neutral and reward-related social odors. The olfactory information left by a rewarded mouse helps other individuals find the reward in a novel environment. Social reward-related olfactory cues change the pattern of social interactions, by increasing following of other animals in the group. Importantly, degree to which the number of followings increases in the presence of reward-related social cues reflects the individual differences in social status within the group. Further, we show that impairing neuronal plasticity in the PL with TIMP-1 results in significantly diminished persistence in seeking for social reward-related olfactory cues, lack of increase in following other animals, and impaired detection of the reward cues in a novel environment.

Presented behavioral paradigm can be further used to study mechanisms of social learning, and in particular social learning strategies^36^. Reward learning, in contrast to threat learning, is more dependent on when and from whom one learns to optimize behavior^37,38^. As searching for reward is an investment, being able to estimate the effort needed to get it and attractiveness of the bounty are important. Thus, the reliability and accessibility of the source of knowledge is key.

We show that olfactory cues from well-known, familiar conspecifics indeed suffice for the transfer of information about the reward. This observation is in line with information theory, according to which in a novel environment animals with no direct experience heavily rely on social cues^17^. Interestingly, we observed that after the two scout mice from the cohort had been patrolling and scent marking the novel environment, the rest of the mice, subsequently moved into the Eco-HAB, preferred drinking from the bottles that had been used more frequently by their peers. This effect was more pronounced in the experimental group, where scout mice had been given access to reward, but also present in the control condition, when scouts had drunk water but, as usually observed in freely behaving mice, more eagerly from one of the presented bottles. Albeit this effect was transient, it shows the value of social information when individual experience is missing^39^. Effects of gaining preference for the things liked by the members of one’s social group are well-documented in humans^39^. Presented experimental paradigm opens new avenues of investigation of such phenomena and enables better understanding of their neural background in well-controlled rodent experiments.

The prefrontal cortex (PFC) is essential for successful navigation through a complex social world both in humans and in other social species, especially the dorsomedial and medial PFC (dmPFC and mPFC) have been implicated in monitoring of emotions and actions in self and others^39^. The functional homolog of these regions in rodents is the dorsal mPFC, including the anterior cingulate cortex and the prelimbic cortex (PL)^40^. We show that disrupting neuroplasticity in the PL with TIMP-1 results in abolished response to social olfactory cues indicating reward. This is in line with the earlier study showing that the PL neurons selectively respond to social olfactory cues^35^.

Further, social learning relies on efficient social communication^36^ and social information processing^38^. We have developed a method of tracking social interactions between individual mice living in a group in the Eco-HAB by tracing movement through the corridors of the system. We observed that social olfactory information about the reward in the environment increased the average number of such interactions, and TIMP-1 release in the PL blocked that increase. Since changes in the followings showed high individual variability they may play a role in social communication. Thus, we looked into the network of interactions within the group and their relationship with social hierarchy.

Social structure in humans can be described by topological variables such as centrality, social distance and betweenness^41^. The most common visualization is the graph with nodes representing individual subjects, edges describing relations between them, and centrality and social distance displayed as their spatial positions^42^. Here we used the number of followings of other mice within the group to analyze the social relationships and show that mice, similarly to primates, form stable social networks. Human studies show that information is spread mostly by so-called information hubs, i.e., humans that have many social connections^43^. Interestingly, our studies show that being a social hub in mice is related to a higher increase in following others when a reward-related social cues appear in the environment. Thus, the Eco-HAB measures we have developed allow for tracking individual variability in social interactions, which seems to be crucial for information spreading.

We show a positive correlation between position in social network and dominance measures. This result suggests that, at least partially, number of followings performed in the Eco-HAB system may be related to hierarchy formation and maintenance. This is also in line with the recent rodent works, which have shown that learning and experience-dependent synaptic plasticity in the PFC are critical for social rank status^22,38,39^. Similarly, when impairing neuronal plasticity in the PL we observed flattening of the social network. The imperfect linear relationship between the traditionally measured hierarchy and followings in the social network suggests that linear model of social hierarchy does not appropriately reflect complexity of the phenomenon. Thus, the detailed relation of social network to hierarchy requires further studies.

Tracking individual differences in social interactions through social networks measured in the Eco-HAB can be very useful for modelling social impairments. In most of the classical tests of sociability randomly chosen pairs of mice are tested, which can increase variability of the results and blur the conclusions. In contrast, longitudinal observation of mice in the Eco-HAB system enables tracking of complex, voluntary and dynamic interactions, thus far exceeding the limits of what can be studied in the conventional approaches to measuring social behavior.

## Figure legends

**S1. Social olfactory cues indicating reward do not change locomotor activity**.

**(A)** Total number of visits performed by the mice from cohort 1.

**S2. Disrupting synaptic plasticity in the prelimbic cortex does not change locomotor activity**.

**(A)** Total number of visits performed by the mice from cohort 2.

**S3. Locomotor activity in experiments on social transfer of information in a novel environment**.

**(A)** No difference in locomotor activity was observed between control (**CTRL-vehicle**) and reward (**REW-vehicle**) groups. In contrast, TIMP1 injection to the PL reduced general activity.

**S4. Observation of the social network in naïve animals** (cohort 6) **shows that it is formed very early in the experiment and remains stable throughout. (A)** Social network of naïve mice housed in the Eco-HAB system without any additional stimulation for 4 days (upper row); the same cohort was then subjected to the injection of TIMP1-carrying NPs into the PL and reintroduced to the Eco-HAB environment for another 4 days (lower row). The social network graphs and the respective horizontal graphs (**B, C**, corresponding to those in E, H of Fig. 4) show a stable level of following during control period (B) and its suppression resulting from the disruption of the neuronal plasticity in the PL (C). As previously described in Fig. 4., patterns of followings between individuals form group’s social network represented as a weighted, directed graph with nodes corresponding to individual mice and edges to interactions between them. Different colors represent the followings of each individual mouse. The numbers next to each node give the PageRank centrality of that node calculated for the weighted, directed graph represented by the adjacency matrix of followings; the PageRank centrality is expressed as a percentage (rounded to the closest integer), characterizing each mouse in relation to a whole social group summing up to 100%. The radius of the colored circle at a given node is proportional to the number of followings performed by the corresponding mouse, while the radius of the dashed circle is proportional to the number of leadings performed by that mouse. The arrows are directed from a follower to a leader; the thickness of an arrow is proportional to the number of followings a given follower performed after a given leader. Horizontal segments (B, C) correspond to the distribution of the number of following events in all pairs of mice within cohort 6. in the subsequent dark phases of the experiment (1-4). Subsequent days of the experiment correspond to respective rows, from bottom to top. Rectangles running from left to right reflect the number of followings ranging from 0 to 40. The intensity of the shading represents the number of occurrences of a given number of followings in all pairs of mice. Additionally, the mean number of followings for all pairs of mice during a given experimental phase, calculated for this distribution, is marked as a red line at the position corresponding to the red scale at the upper edge of the plot.

**S5. The number of followings performed by individual mice corresponds with their social status**.

**(A)** Schematic of the experimental design. First, followings were measured in the Eco-HAB, then the U-tube dominance test was performed. (**B)** Positive correlation between the position within the social network, defined by the number of followings performed in the Eco-HAB, and dominance hierarchy as defined by the U-tube test. The dominant mice were the ones following others the most.

## Materials and Methods

### Subjects

Animals were treated according to the ethical standards of the European Union (directive no.2010/63/UE) and respective Polish regulations. All the experiments were preapproved by the Local Ethics Committee no. 1 in Warsaw, Poland. C57BL/6 male mice were bred in the Animal House of Nencki Institute of Experimental Biology, Polish Academy of Sciences or Mossakowski Medical Research Centre, Polish Academy of Sciences. The animals entered the experiments when 2-3 month old. They were littermates derived from several breeding pairs. The mice were transferred to the animal room at least 2 weeks before the experiments started and put in the groups of 12-15 in one cage (56 cm x 34 cm x 20cm) with enriched environment (tubes, shelters, nesting materials). They were kept under 12h/12h light-dark cycle. The cages were cleaned once per week.

### RFID tagging

Glass coated RFID microtransponders (9.5 mm - length and 2.2 mm - diameter, RFIP Ltd) were sterilized in 70% ethanol, dried on a paper towel, loaded into the syringes and injected subcutaneously into the subjects anesthetized with isoflurane. Each transponder had a unique number that can be registered by the Eco-HAB antennas when animals pass through its corridors. After injections mice were put together into one home cage and moved back to the experimental room. On the next day the presence and the position of the transporters under animals’ skin was additionally verified.

### Poly(DL-lactide-co-glycolide) nanoparticles containing TIMP-1 or BSA

To gradually release TIMP-1 in the PL of the tested animals we used poly(DL-lactide-co-glycolide, PLGA) nanoparticles (NP) loaded with the protein (in the control condition bovine serum albumin, BSA, Sigma-Aldrich). The NPs were prepared according to the protocol described by Chaturvedi et al.^29^. Briefly, NPs were prepared in the process of multiple emulsifications and evaporations (MW 45.000–75.000, copolymer ratio: 50:50, Sigma-Aldrich). 100 mg PLGA was dissolved in 5 ml dichloromethane and 4ml of dimethyl tartaric acid (Signa-Aldrich). In the next step, 1 mg of TIMP-1 or BSA was dissolved in 500 μl of MiliQ water. The protein solution was mixed with dichloromethane containing PLGA, sonicated and emulsified in 1% polyvinyl alcohol (on average MW 30.000–70.000, Sigma-Aldrich). Additionally, FITC was added to easily localize place of NP’s delivery in the brain. Subsequently, the solution was stirred at room temperature overnight to evaporate dichloromethane. Next, the NPs were centrifuged at 10.000 x g, washed three times with MiliQ, dissolved in PBS, and stored at 4°C.

### Stereotaxic surgeries

All tools were sterilized in 70% ethanol before the surgical procedures. Mice were anesthetized by isoflurane inhalation (started at 5% and reduced to 2-1,5% of isoflurane) and placed in a stereotaxic apparatus (Kopf Instruments) on a heating pad (37.8°C). The mice were subcutaneously injected with analgesic (Butamidor, Richter, 1:20 in saline, 2.5 ml/kg) and reflexes were checked to ensure absence of pain. To protect animals’ eyes from drying we used a moisturizing gel (Carbonerum, Vidisic). Ear bars were put into place and scalp was shaved. The skin on the skull was cut to expose bregma. Nanofil 35G needles were used to bilaterally inject NPs into the PL (coordinates: AP +1.8 mm, LM +/- 0.92 mm, DV -1.67 mm, at 20° angle). The delivery was controlled by the Micro Syringe Pump (World Precision Instruments, 500 nl of total volume, 100 nl/min). To let the NPs diffuse the needle was left in the brain for additional 5 min after the injection. Afterwards, the incision was sutured (Dafilon, C0935204) and lubricated with the analgesic lignocainum hydrochloricum (10 mg, Polfa). The mice received subcutaneous injections of anti-inflammatory medication (Tolfedine, Vetoquinol, 4 mg/kg) and a wide-spectrum antibiotic (Baytril 2.5%, Bayer, 1:3 in saline, 5 ml/kg). Then mice were placed in cages warmed with a heating pad and singly-housed for the next 5 days to allow for full recovery.

### Perfusions and verification of TIMP-1 injections

After the end of behavioral testing, mice injected with the NPs releasing TIMP-1 or BSA were anesthetized with intraperitoneal injection of sodium pentobarbital (100 mg/kg, dissolved in PBS) and transcardially perfused with 80 ml of ice-cold PBS followed by 60 ml of 4% PFA in PBS (4°C). The brains were isolated and placed overnight in 4% PFA in PBS (4°C). Then, the brains were moved to 30% sucrose solution in PBS for 2-3 days (4°C) for cryoprotection. Afterwards, the brains were cut on a cryostat into 50 μm-thick coronal slices. The slices were then washed in PBS, placed on the microscope glass slides and fluorescence of FITC encapsulated in NPs was imaged under the Nikon Eclipse Ni microscope.

### Eco-HAB system

The Eco-HAB is a fully automated, open source system for testing social behavior in group-housed mice living under the 12/12h dark/light cycle^30^. The system comprises of 4 polycarbonate compartments (30cm x 30cm x 18cm) connected with tube-shaped corridors (inner diameter 36mm, outer diameter 40mm), and covered with stainless steel grid lid. In 2 out of 4 compartments mice have access to food and water (*ad libitum*); the other 2 compartments have a separated space for presentation of the olfactory stimuli (in a corner, behind a perforated partition) or putting additional bottles. Access to all compartments, olfactory stimuli and additional bottles is unrestricted and voluntary. All housing elements can be autoclaved and disinfected with 70% alcohol. To record movement of the animals within the Eco-HAB the corridors are equipped with circular RFID antennas registering transponders’ numbers. The data from the antennas are collected by a dedicated electronic system, which sends them to the computer. Eco-HAB measures were computed with pyEcoHAB library (https://github.com/Neuroinflab/pyEcoHAB). The following behavioral measures were analyzed. **In a familiar environment approach to social odor** was measured as a proportion of visits to the compartments with olfactory stimuli (reward:neutral or neutral:neutral, depending on the experiment) divided by the same proportion from the preceding dark phase (when no olfactory stimuli were present in the compartments). **In the experiments performed in a novel environment preference to the bottles** was measured as a relative time spent on drinking from the preferred bottle in relation to the non-preferred bottle in 1^st^ hour of the experiment. **Persistence** was defined as a proportion of visits to the compartments where odors were presented (reward:neutral or neutral:neutral) during the last 6 hours of the testing phase, divided by the same proportion from the first 6 hours of the testing phase. This measure shows persistence in seeking for social information about the reward. In the experiments performed in the novel environment persistence was calculated similarly to what was previously described, but the time spent on drinking from the additional bottles was divided by the time spent in chambers where the bottles were presented. **Followings** were measured based on the number of events when mice followed one another in the corridors of the Eco-HAB system. Specifically, following was defined as an event when one mouse entered a corridor, followed by another mouse before the first left the tube, and when both mice left the corridor in the same order and in the same direction. For the purpose of between-group comparisons to control for the individual variability in locomotor activity we summed all the following events of each mouse and divided by its activity (total number of its visits) during the analyzed time bin. For the details of the implemented algorithm please refer to https://github.com/Neuroinflab/pyEcoHAB. For the analysis of within-cohort changes in following patterns presented in Fig 4 raw values were used.

### Testing behavioral response to social olfactory cues indicating reward presented in a familiar environment

13 mice (cohort 1) were put into the Eco-HAB system at the beginning of the dark phase. Please note that one mouse had to be excluded from experiments due to health problems with fore tooth overgrowth that might have affected its social behavior. Animals were adapted to the testing environment for 48 hours (Adaptation phase). Then, 2 randomly chosen mice were separated and housed in the individual cages (17 cm x 23 cm x 13 cm) for the next 24h. They were offered either tap water or 10% sucrose solution, food was freely available (Isolation phase). Bedding soaked with the scents of the separated mice was used as social olfactory cues; for the next 24h it was put behind the perforated partitions in the Eco-HAB system to avoid spreading it throughout the cages while maintaining unrestricted sniffing. The cohort was tested twice, in the first (control-CTRL) trial both separated mice had access to tap water, in the second trial (experimental-REW) the same mice were isolated and one of them had access to highly rewarding 10% sucrose solution, while the other to tap water. The design of the experiment with TIMP-1 released into the PL was performed in accordance to the same protocol. Similarly, the mice (cohort 2, n=13) were tested twice, before and after brain manipulation (5 days after NPs injection). As previously described, in both trials, the same pair of mice was isolated to produce social olfactory cues.

### Testing behavioral response to social olfactory cues indicating reward presented in a novel environment

#### Social olfactory information in a new environment

The mice were injected with NPs containing TIMP1 or BSA (vehicle) 5 days before the start of the experimental procedures. We tested 3 cohorts of mice: CTRL-vehicle (cohort 3, n = 12, one mouse had to be excluded from the analysis because it had not drunk), REW-vehicle (cohort 4, n = 12) and REW-NP-TIMP1 (cohort 5, n=12, 2 mice died after the surgeries). Two opposite Eco-HAB compartments offered access to the additional bottles through a short (8cm long) tube, equipped with an RFID antenna to register drinking time of each mouse. As in the previous experiments, the mice were put into the Eco-HAB system (Eco-HAB I) at the beginning of the dark phase. They were adapting to the environment for the next 72 hours (Adaptation phase). Then, 2 randomly chosen mice were moved into a new, clean Eco-HAB system (Eco-HAB II), in which the only bottles accessible were the ones with the antennas monitoring drinking behavior. In the reward (REW) trials the separated mice had access to tap water in one compartment and to 10% sucrose solution in the opposite compartment of the Eco-HAB II. In the control (CTRL) trial both bottles were filled with tap water. After 24 h of the Separation phase, the two scout mice were removed, the bottles replaced with the ones containing fresh tap water. Then the rest of the group, which until now had inhabited the original Eco-HAB I, was transferred to Eco-HAB II for 12 h (Testing phase).

#### Longitudinal observation of social structure formation in the Eco-HAB

To observe how social structure is formed and how it is affected by TIMP-1 release in the PL we tested voluntary behavior of mice in the longitudinal experiments, in which animals inhabited Eco-HAB, without any additional changes to the testing environment. The mice (cohort 6, n=15) were placed into the Eco-HAB system and observed for 4 days. Next, they were subjected to the stereotaxic injections with TIMP-1 loaded NPs. After recovery the mice were placed back into the Eco-HAB and their behavior was measured for another 4 days.

#### Dominance tests

To assess the relationship between social structure and dominance hierarchy we tested the same subjects (cohort 7, n=12) in the Eco-HAB and U-tube tests. Following 10-day-long observation of the social structure in the Eco-HAB system, the mice were subjected to the U-tube dominance test. Mice were placed in single cages and tested in all possible pairwise combinations. The mice from the currently tested pair were placed at the opposite ends of the U-shaped tube (1m length, 42 mm diameter) and allowed to interact. When one mouse pushed the other out it won a given bout. We tested all pairs and calculated the number of wins for each subject as a measure of dominance.

#### Technical reports

As all Eco-HAB data are recorded automatically we verified its integrity to ensure there were no corrupted segments or other problems that might have affected the results. In rare situations of brief antennas’ malfunctions we were able to detect such situations and present the results in supplemental reports (Fig S6 – Fig S12). It is noteworthy, that applied algorithms include heuristics that account for an occasional missed reading of an antenna, to mitigate the impact of such problems on the results. As a rule, if the percentage of antenna errors exceeds the level of 5%, the experiment is considered technically invalid and data is not analyzed. No such problem occurred in the presented studies.

Fig S6. Technical report – cohort 1: reward information sharing experiments

Antennas mismatches did not exceed the level of 5% and the experiment was considered valid for further analysis

Fig S7. Technical report – cohort 2: reward information sharing after TIMP-1 experiments Antennas mismatches did not exceed the level of 5% and the experiment was considered valid for further analysis

Fig S8. Technical report – cohorts 3: reward information sharing experiments in novel environment, control group.

Antennas mismatches did not exceed the level of 5% and the experiment was considered valid for further analysis

Fig S9. Technical report – cohorts 4: reward information sharing experiments in novel environment, vehicle-treated group

Antennas mismatches did not exceed the level of 5% and the experiment was considered valid for further analysis

Fig S10. Technical report – cohort 5: reward information sharing experiments in novel environment, TIMP1-treated group

Antennas mismatches did not exceed the level of 5% and the experiment was considered valid for further analysis

Fig S11. Technical report – cohort 6: longitudinal observation of the social network

Antennas mismatches did not exceed the level of 5% and the experiment was considered valid for further analysis

Fig S12. Technical report – cohort 7: experiments on the relationship of the social network with dominance hierarchy

Antennas mismatches did not exceed the level of 5% and the experiment was considered valid for further analysis

### Statistical analyzes

For statistical analyses we used GraphPad Prism7 software. The normality of data distributions was assessed with the D’Agostino-Pearson omnibus normality test. Data sets that passed the normality tests were further analyzed using Student’s t-test for independent or paired samples, depending on the particular type of comparison. For the data sets that did not pass the criterion of normality required for performing parametric analyses, the Mann-Whitney or Wilcoxon matched-pairs signed-rank tests were used. For comparisons of data with the theoretical value (no change level) we used one sample t-test for parametric data. Correlation was calculated by Pearson correlation test. To assess the significance of each mouse as an element of the social network, we calculated PageRank centrality for the inverted weighted directed graph of followings or, equivalently, for the weighted directed graph of leading, in which each directed edge carries a weight equal to the number of respective events^44^. Calculations were performed with the use of Wolfram Mathematica. The weights are given in percent and rounded to the nearest integer number. The criterion for statistical significance was a probability level of p<0.05. Statistical details for all experiments are described in Table S1.

## Authors contributions

Concept and design M.W., A.P., E.K.; data acquisition M.W., J.B., M.R.W., L.K., M.C.; analysis and interpretation of data M.W., J.B., M.R.W., J.J.S., D.K.W., L.K., K.T. A.P., E.K.; drafting and revising the article M.W., L.M., D.K.W., K.T., A.P., E.K.

## Bibliography

1. Kaplan, H. S., Hooper, P. L. & Gurven, M. The evolutionary and ecological roots of human social organization. Philos Trans R Soc Lond B Biol Sci 364, 3289–3299 (2009).

2. von Rueden, C. R., Redhead, D., O’Gorman, R., Kaplan, H. & Gurven, M. The dynamics of men’s cooperation and social status in a small-scale society. Proc. R. Soc. B. 286, 20191367 (2019).

3. Brent, L. J. N., Chang, S. W. C., Gariépy, J.-F. & Platt, M. L. The neuroethology of friendship: Neuroethology of friendship. Ann. N.Y. Acad. Sci. 1316, 1–17 (2014).

4. Pérez-Escudero, A. & de Polavieja, G. G. Adversity magnifies the importance of social information in decision-making. J R Soc Interface 14, (2017).

5. Reader, S. M. Animal social learning: associations and adaptations. F1000Res 5, 2120 (2016).

6. Márquez, C., Rennie, S. M., Costa, D. F. & Moita, M. A. Prosocial Choice in Rats Depends on Food-Seeking Behavior Displayed by Recipients. Current Biology 25, 1736–1745 (2015).

7. Langford, D. J. et al. Social modulation of pain as evidence for empathy in mice. Science 312, 1967–1970 (2006).

8. Han, Y. et al. Bidirectional cingulate-dependent danger information transfer across rats. PLoS Biol 17, (2019).

9. Jeon, D. et al. Observational fear learning involves affective pain system and Cav1.2 Ca2+ channels in ACC. Nat Neurosci 13, 482–488 (2010).

10. Lloyd, J. A. Social structure and reproduction in two freelygrowing populations of house mice (Mus musculus L.). Animal Behaviour 23, 413–424 (1975).

11. Beny-Shefer, Y. et al. Nucleus Accumbens Dopamine Signaling Regulates Sexual Preference for Females in Male Mice. Cell Reports 21, 3079–3088 (2017).

12. Arakawa, H., Blanchard, D. C., Arakawa, K., Dunlap, C. & Blanchard, R. J. Scent marking behavior as an odorant communication in mice. Neuroscience & Biobehavioral Reviews 32, 1236–1248 (2008).

13. Li, Y. et al. Lrfn2-Mutant Mice Display Suppressed Synaptic Plasticity and Inhibitory Synapse Development and Abnormal Social Communication and Startle Response. J. Neurosci. 38, 5872–5887 (2018).

14. Sprott, R. L. “Fear Communication” via Odor in Inbred Mice. Psychol Rep 25, 263–268 (1969).

15. Hurst, J. L. Urine marking in populations of wild house mice Mus domesticus rutty. I. Communication between males. Animal Behaviour 40, 209–222 (1990).

16. Scott, J. W. & Pfaff, D. W. Behavioral and electrophysiological responses of female mice to male urine odors. Physiology & Behavior 5, 407–411 (1970).

17. Duboscq, J., Romano, V., MacIntosh, A. & Sueur, C. Social Information Transmission in Animals: Lessons from Studies of Diffusion. Front. Psychol. 7, (2016).

18. Kondrakiewicz, K., Kostecki, M., Szadzińska, W. & Knapska, E. Ecological validity of social interaction tests in rats and mice. Genes, Brain and Behavior 18, e12525 (2019).

19. Bicks, L. K., Koike, H., Akbarian, S. & Morishita, H. Prefrontal Cortex and Social Cognition in Mouse and Man. Front. Psychol. 6, (2015).

20. Demolliens, M., Isbaine, F., Takerkart, S., Huguet, P. & Boussaoud, D. Social and asocial prefrontal cortex neurons: a new look at social facilitation and the social brain. Social Cognitive and Affective Neuroscience 12, 1241–1248 (2017).

21. Ito, W. & Morozov, A. Prefrontal-amygdala plasticity enabled by observational fear. Neuropsychopharmacol. 44, 1778–1787 (2019).

22. Wang, F. et al. Bidirectional Control of Social Hierarchy by Synaptic Efficacy in Medial Prefrontal Cortex. Science 334, 693–697 (2011).

23. Loureiro, M. et al. Social transmission of food safety depends on synaptic plasticity in the prefrontal cortex. Science 364, 991–995 (2019).

24. Attwood, B. K. et al. Neuropsin cleaves EphB2 in the amygdala to control anxiety. Nature 473, 372–375 (2011).

25. Knapska, E. et al. Reward Learning Requires Activity of Matrix Metalloproteinase-9 in the Central Amygdala. Journal of Neuroscience 33, 14591–14600 (2013).

26. Okulski, P. et al. TIMP-1 Abolishes MMP-9-Dependent Long-lasting Long-term Potentiation in the Prefrontal Cortex. Biological Psychiatry 62, 359–362 (2007).

27. Bach, D. R., Tzovara, A. & Vunder, J. Blocking human fear memory with the matrix metalloproteinase inhibitor doxycycline. Molecular Psychiatry 23, 1584–1589 (2018).

28. Bach, D. R., Näf, M., Deutschmann, M., Tyagarajan, S. K. & Quednow, B. B. Threat Memory Reminder Under Matrix Metalloproteinase 9 Inhibitor Doxycycline Globally Reduces Subsequent Memory Plasticity. J. Neurosci. 39, 9424–9434 (2019).

29. Chaturvedi, M., Molino, Y., Bojja, S., Khrestchatisky, M. & Kaczmarek, L. Tissue inhibitor of matrix metalloproteinases-1 loaded poly(lactic-co-glycolic acid) nanoparticles for delivery across the blood–brain barrier. IJN 575 (2014) doi:10.2147/IJN.S54750.

30. Puścian, A. et al. Eco-HAB as a fully automated and ecologically relevant assessment of social impairments in mouse models of autism. eLife 5, e19532 (2016).

31. Łopucki, R. & Schimrosczyk, D. Recognition of interspecific familiar versus unfamiliar odours among bank voles and yellow-necked mice. Acta Theriologica 48, 167–176 (2003).

32. Lee, R. K.-W., Hoang, T.-A. & Lim, E.-P. Discovering Hidden Topical Hubs and Authorities Across Multiple Online Social Networks. IEEE Transactions on Knowledge and Data Engineering 33, 70–84 (2021).

33. Fuxjager, M. J., Knaebe, B. & Marler, C. A. A single testosterone pulse rapidly reduces urinary marking behaviour in subordinate, but not dominant, white-footed mice. Animal Behaviour 100, 8–14 (2015).

34. Kleshchev, M. A. & Osadchuk, L. V. Social domination and reproductive success in male laboratory mice (Mus musculus). J Evol Biochem Phys 50, 227–233 (2014).

35. Hayashi, S. Development and diversity of social structure in male mice. J. Ethol. 11, 77–82 (1993).

36. Olsson, A., Knapska, E. & Lindström, B. The neural and computational systems of social learning. Nature Reviews Neuroscience 21, 197–212 (2020).

37. Kendal, R. L. et al. Social Learning Strategies: Bridge-Building between Fields. Trends in Cognitive Sciences 22, 651–665 (2018).

38. Parkinson, C., Kleinbaum, A. M. & Wheatley, T. Spontaneous neural encoding of social network position. Nat Hum Behav 1, 0072 (2017).

39. Bandura, A. Social Learning Theory. (Prentice Hall, 1977).

40. Wang, F. et al. Bidirectional Control of Social Hierarchy by Synaptic Efficacy in Medial Prefrontal Cortex. Science 334, 693–697 (2011).

41. Wasserman, S., Urbana-Champaign), S. (University of I. W. & Faust, K. Social Network Analysis: Methods and Applications. (Cambridge University Press, 1994).

42. Bonacich, P. Power and Centrality: A Family of Measures. American Journal of Sociology 92, 1170–1182 (1987).

43. Lee, R. K.-W., Hoang, T.-A. & Lim, E.-P. Discovering Hidden Topical Hubs and Authorities Across Multiple Online Social Networks. IEEE Trans. Knowl. Data Eng. 1–1 (2019) doi:10.1109/TKDE.2019.2922962.

44. Brin, S. & Page, L. The anatomy of a large-scale hypertextual Web search engine. Computer Networks and ISDN Systems 30, 107–117 (1998).

