## Supplementary figures and images for "Social learning about rewards – how information from others helps to adapt to changing environment"

### Fig S1

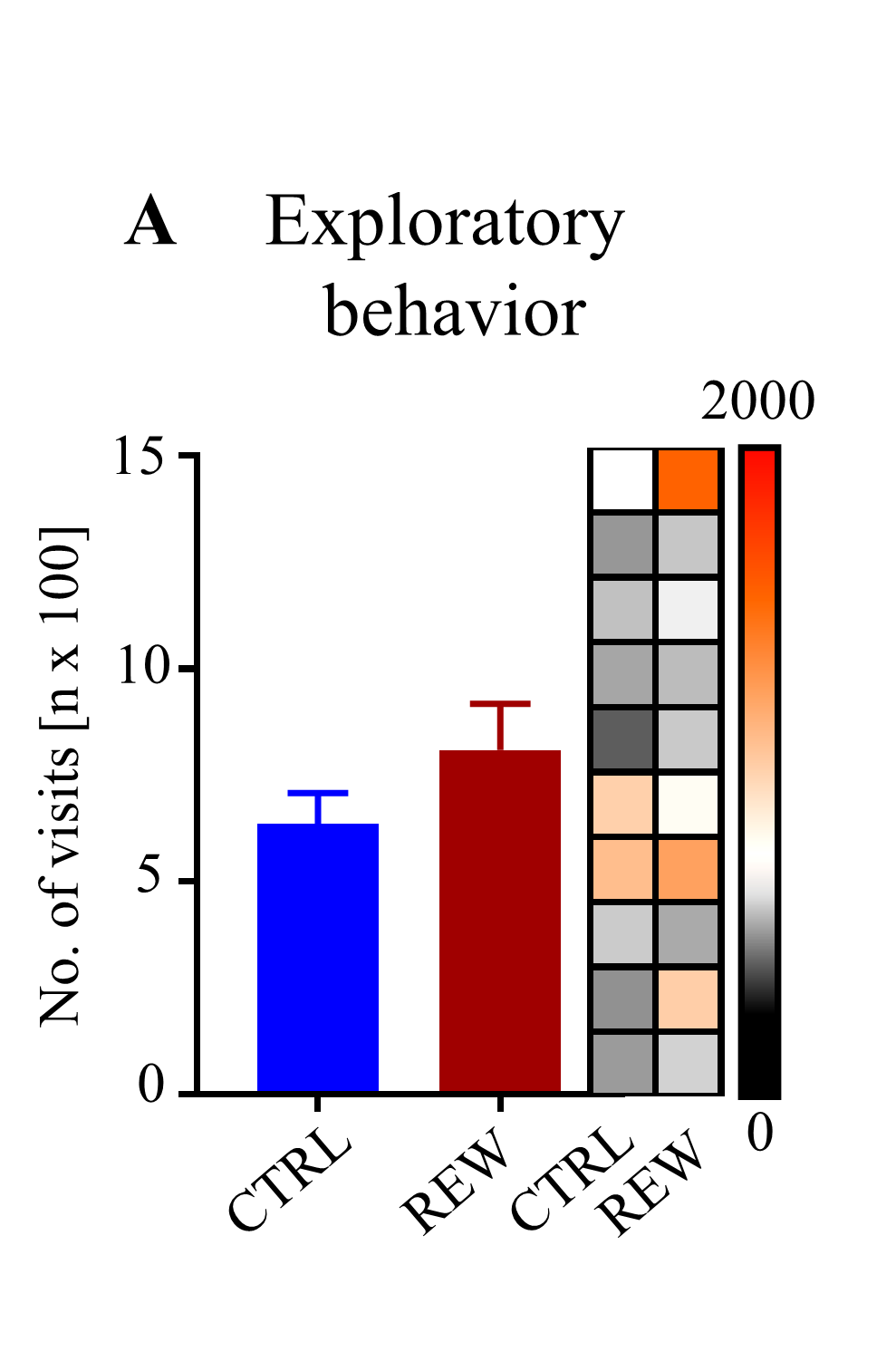

### Fig S2

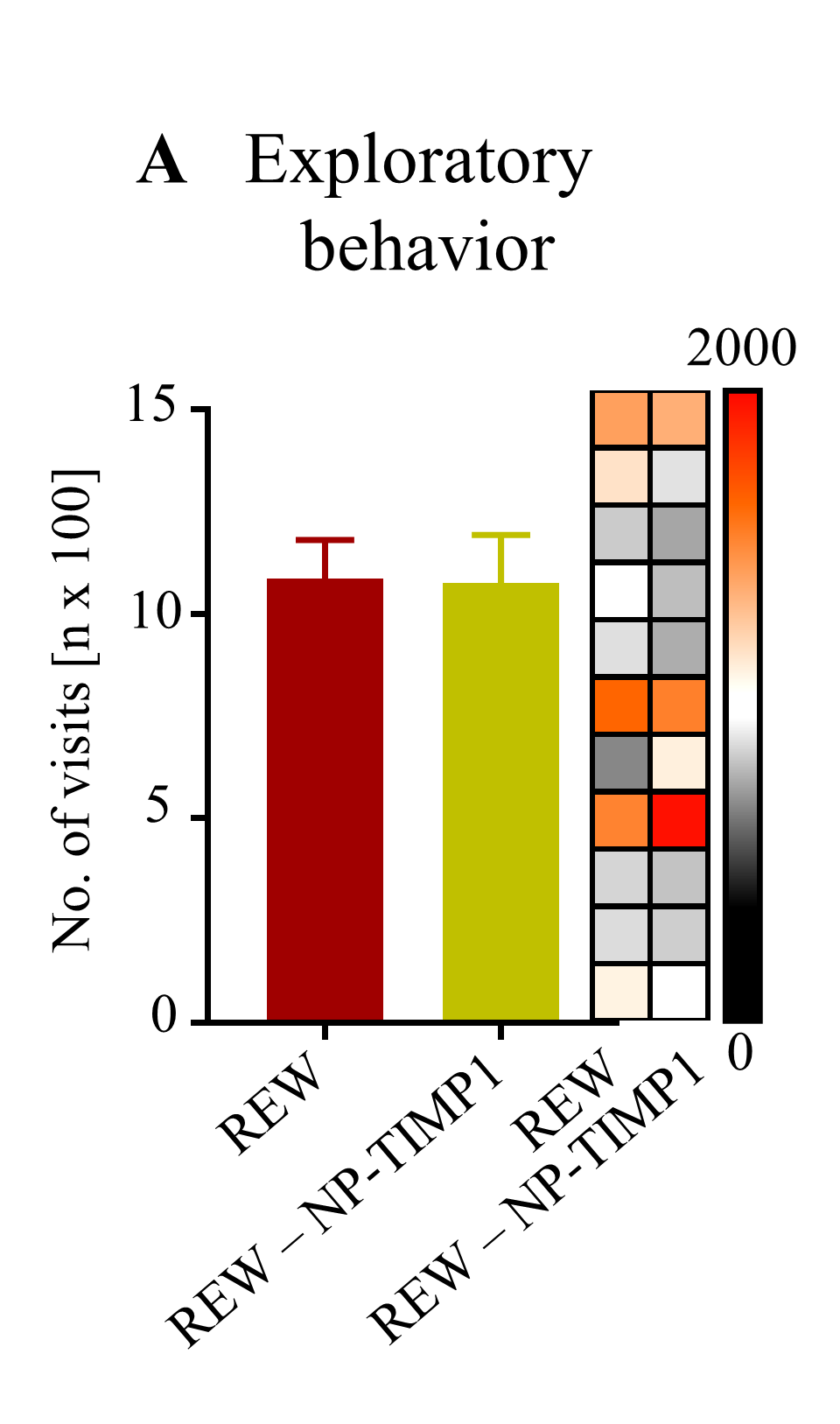

### Fig S3

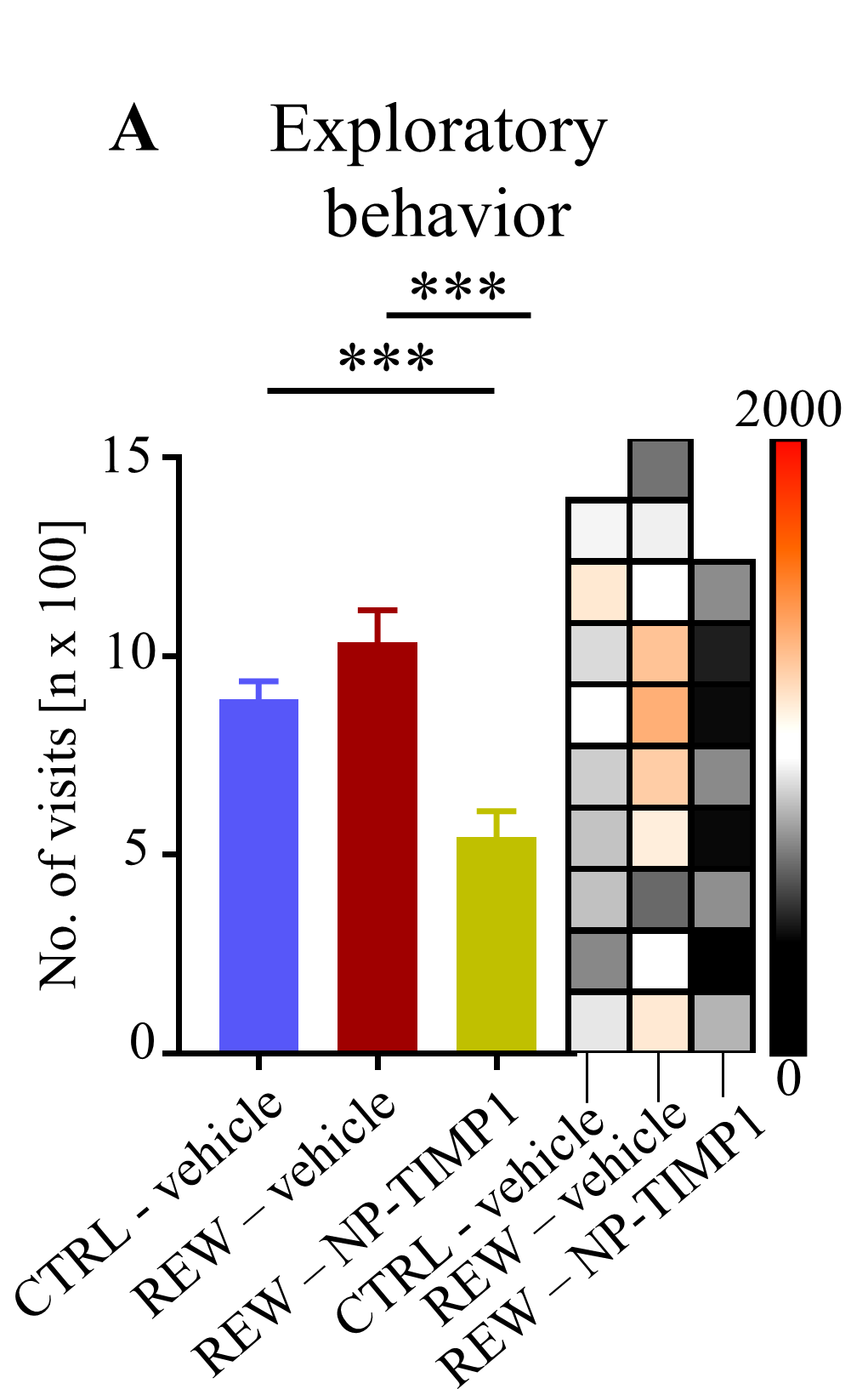

### Fig S4

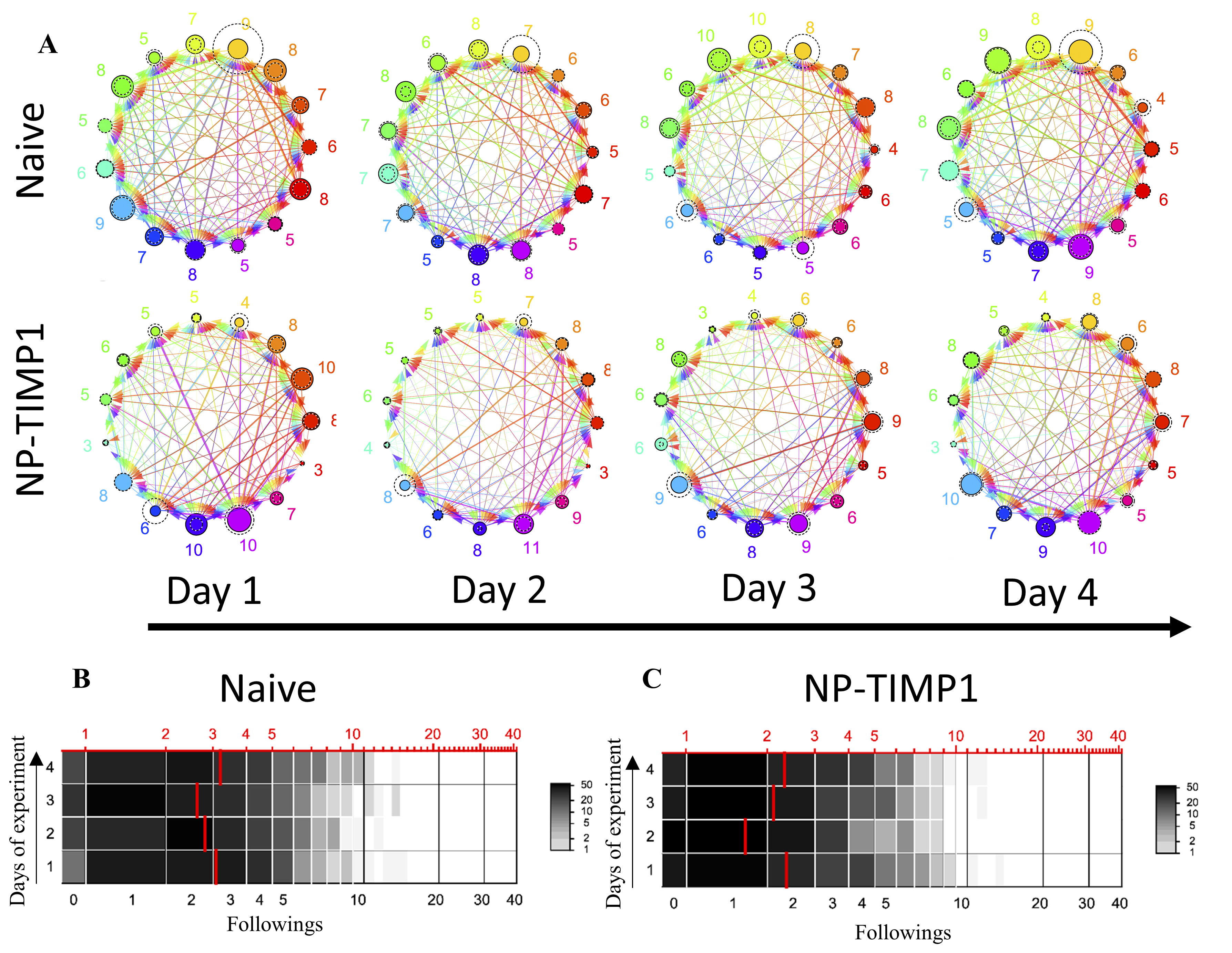

### Fig S5

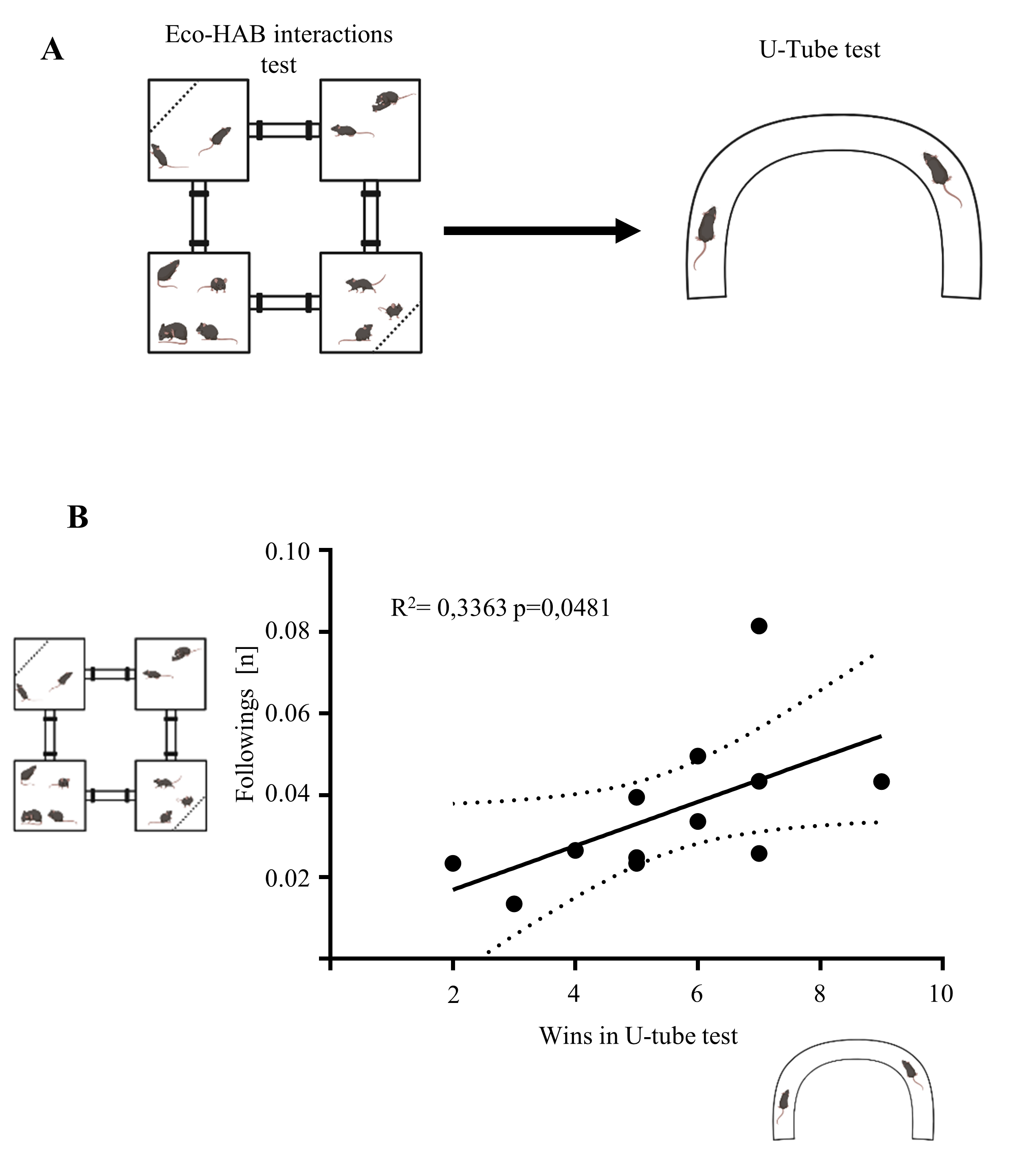

### Fig S6

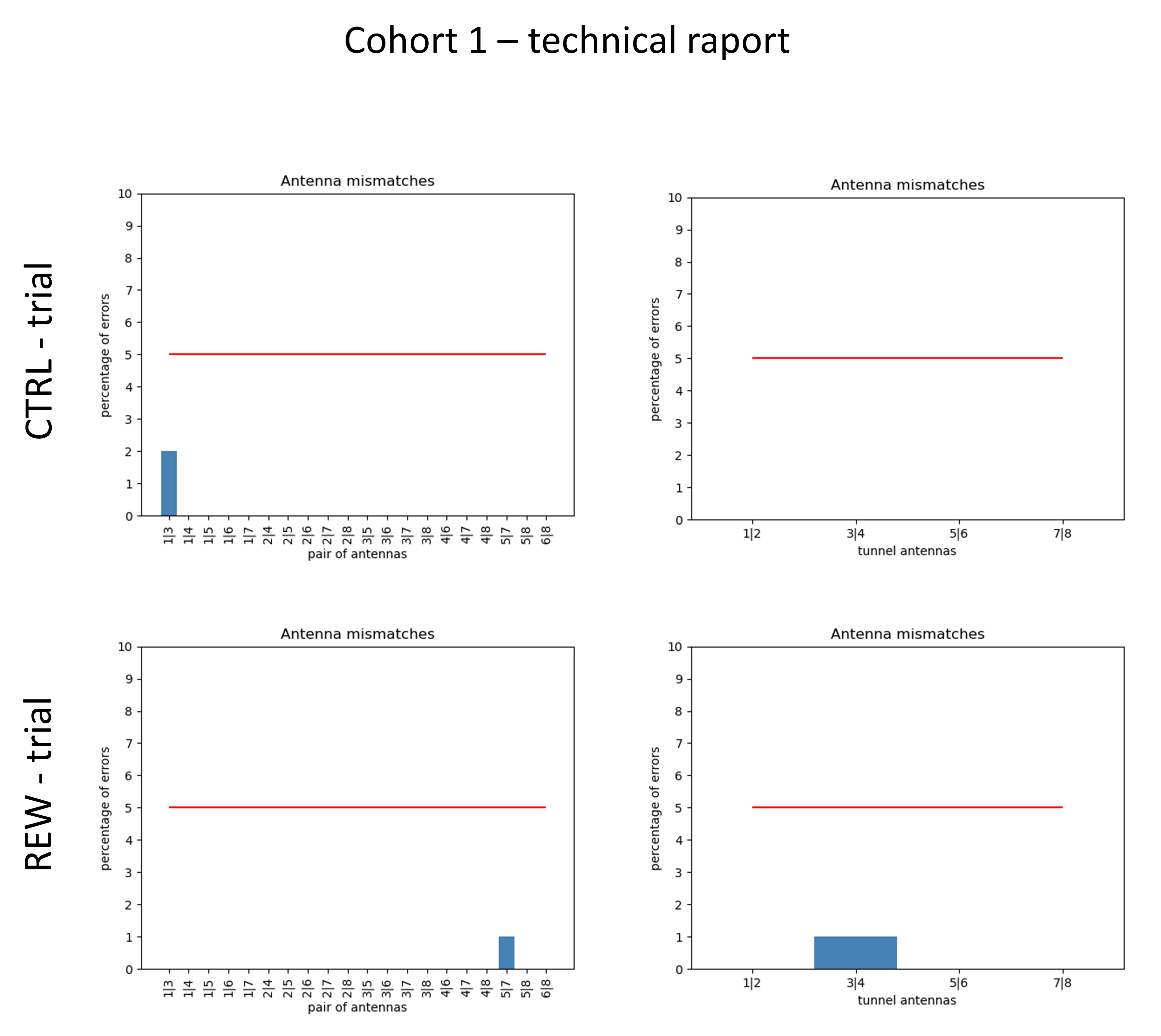

### Fig S7

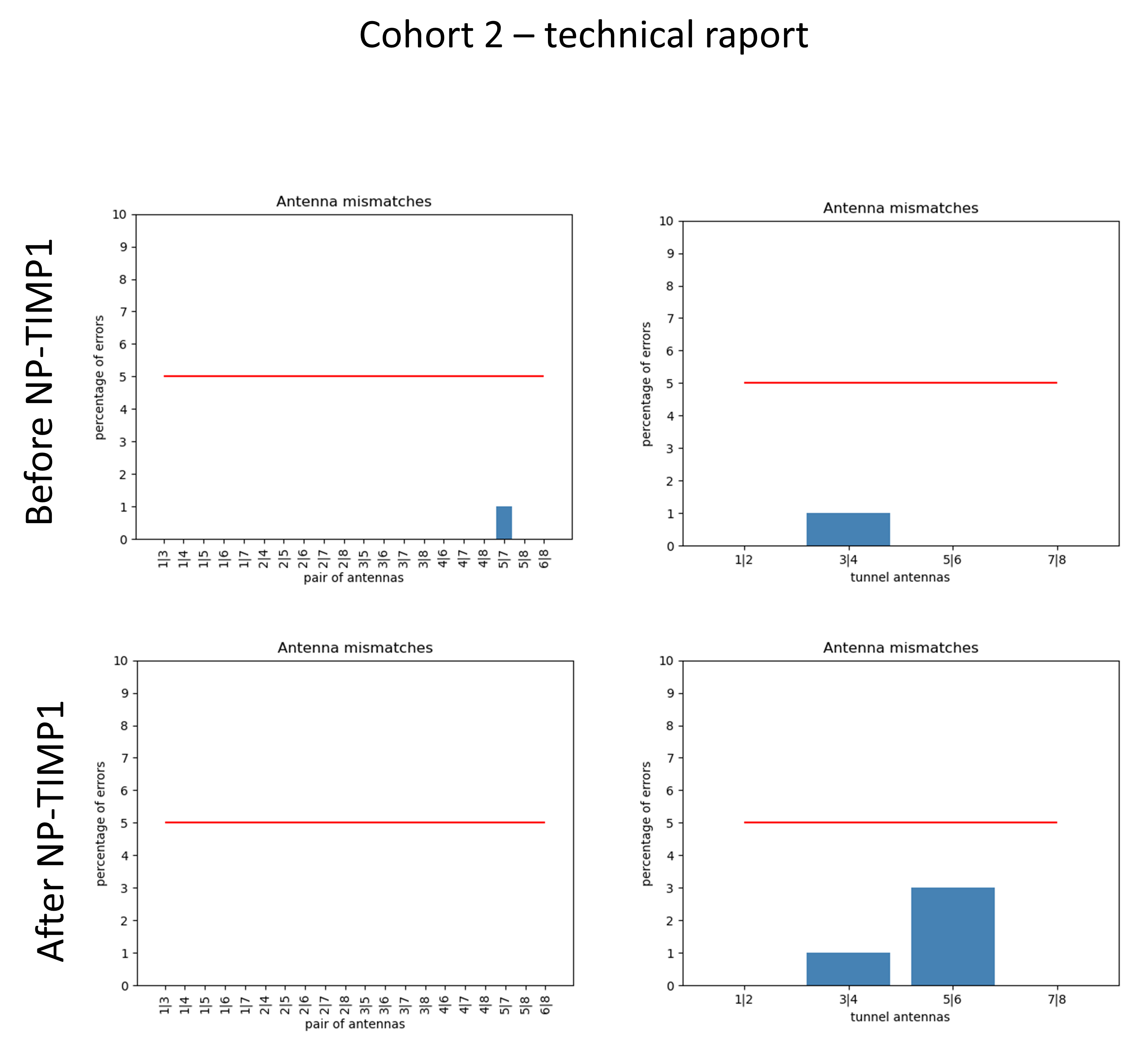

### Fig S8

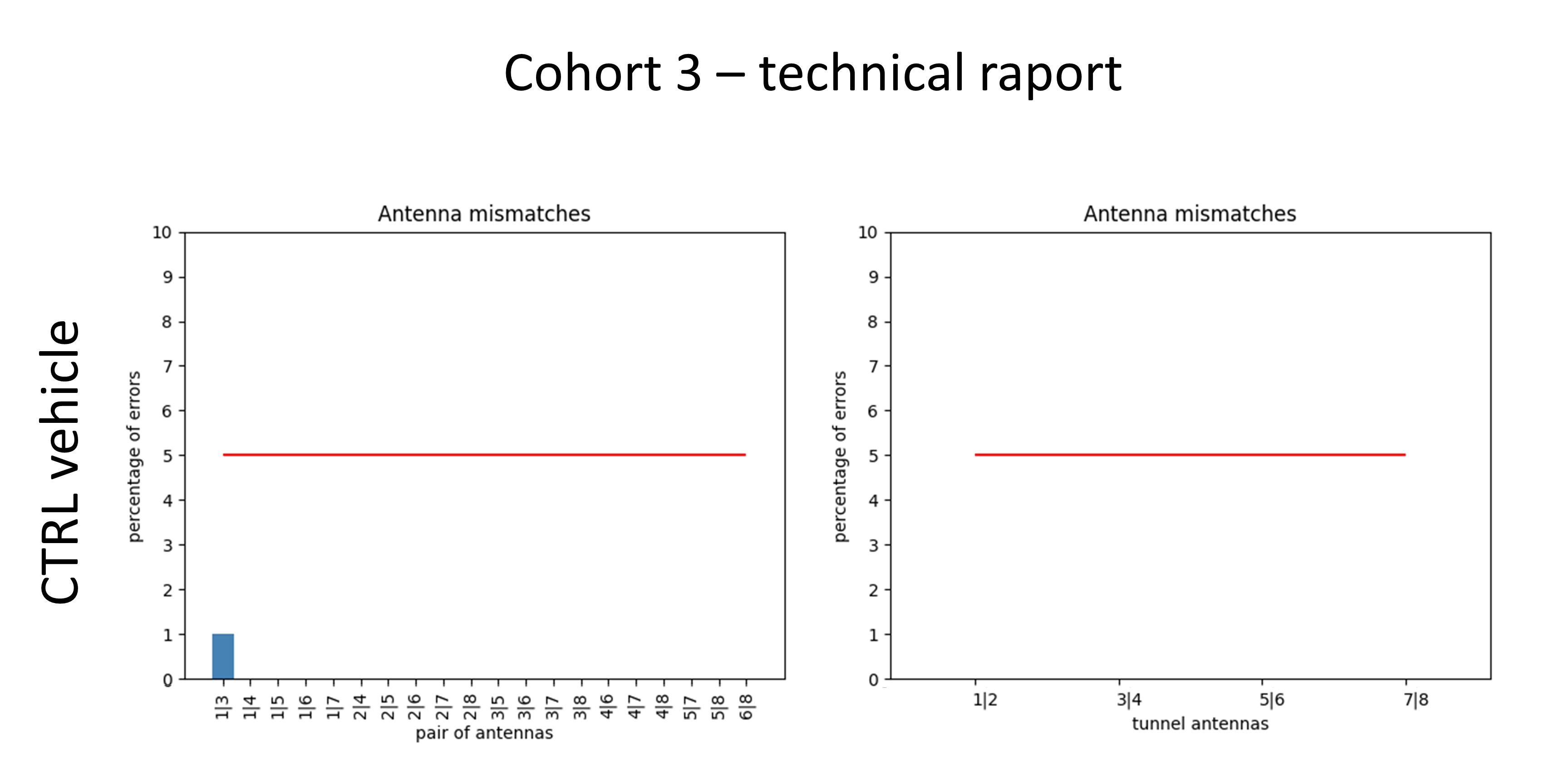

### Fig S9

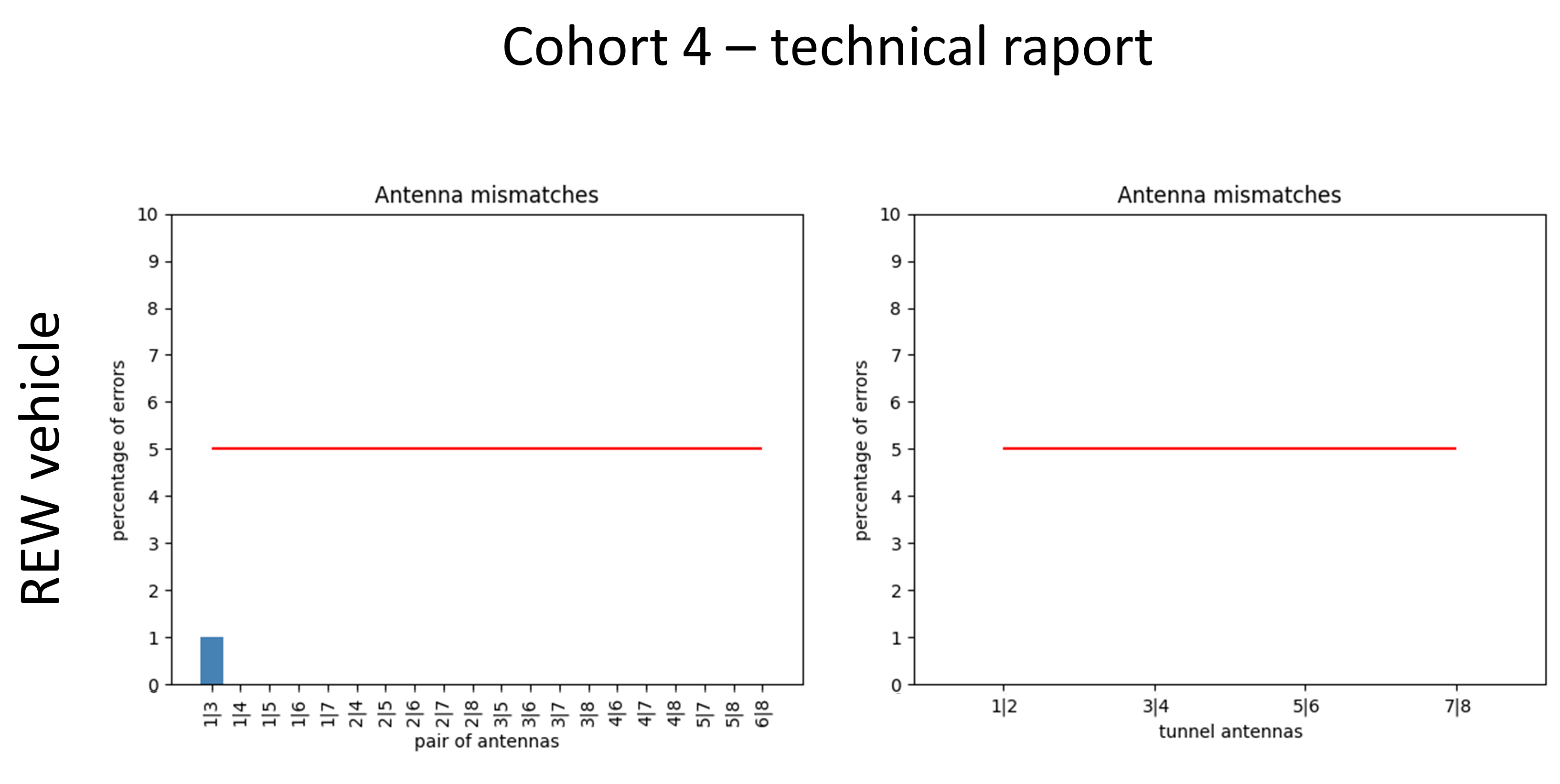

### Fig S10

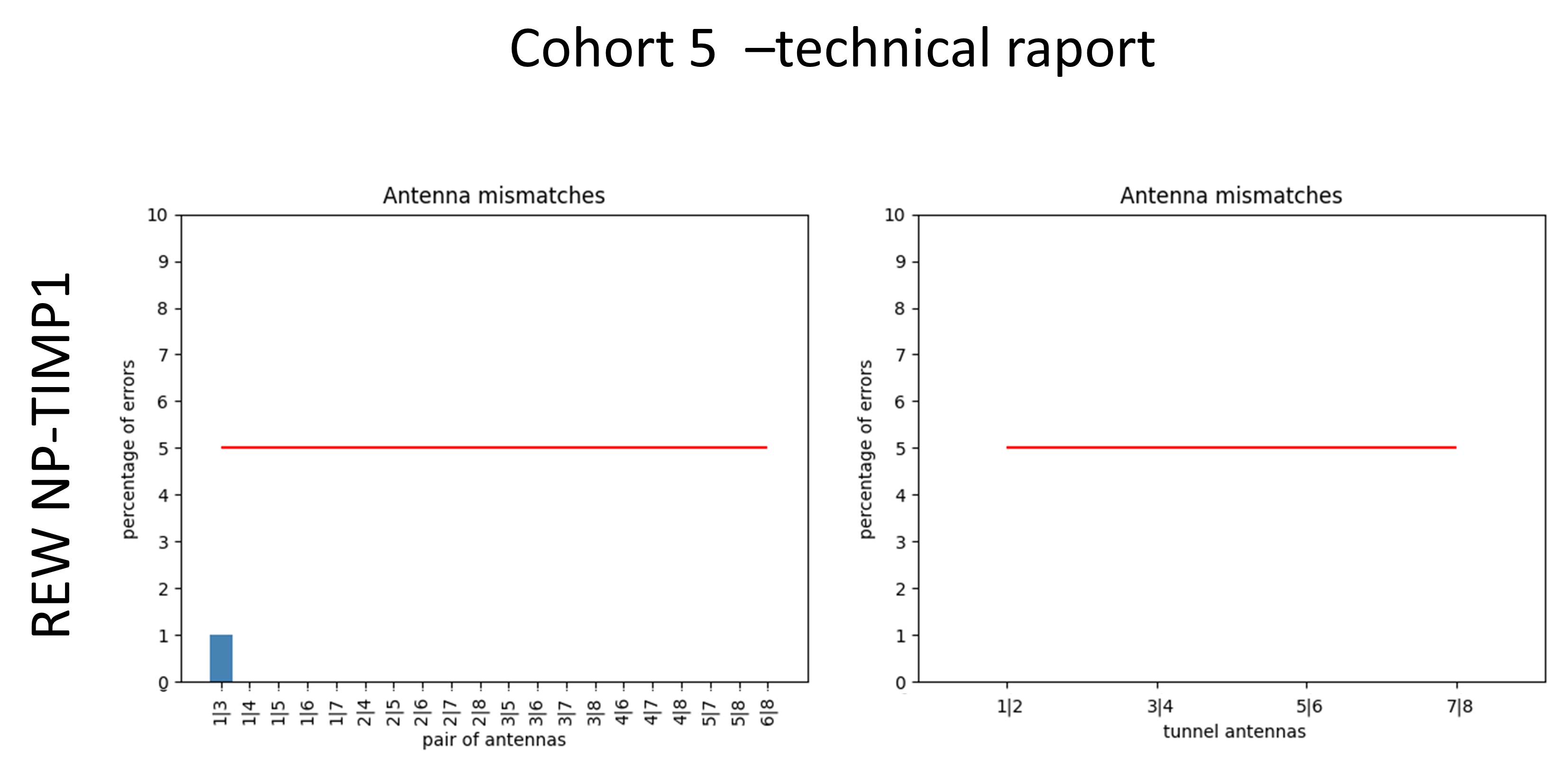

### Fig S11

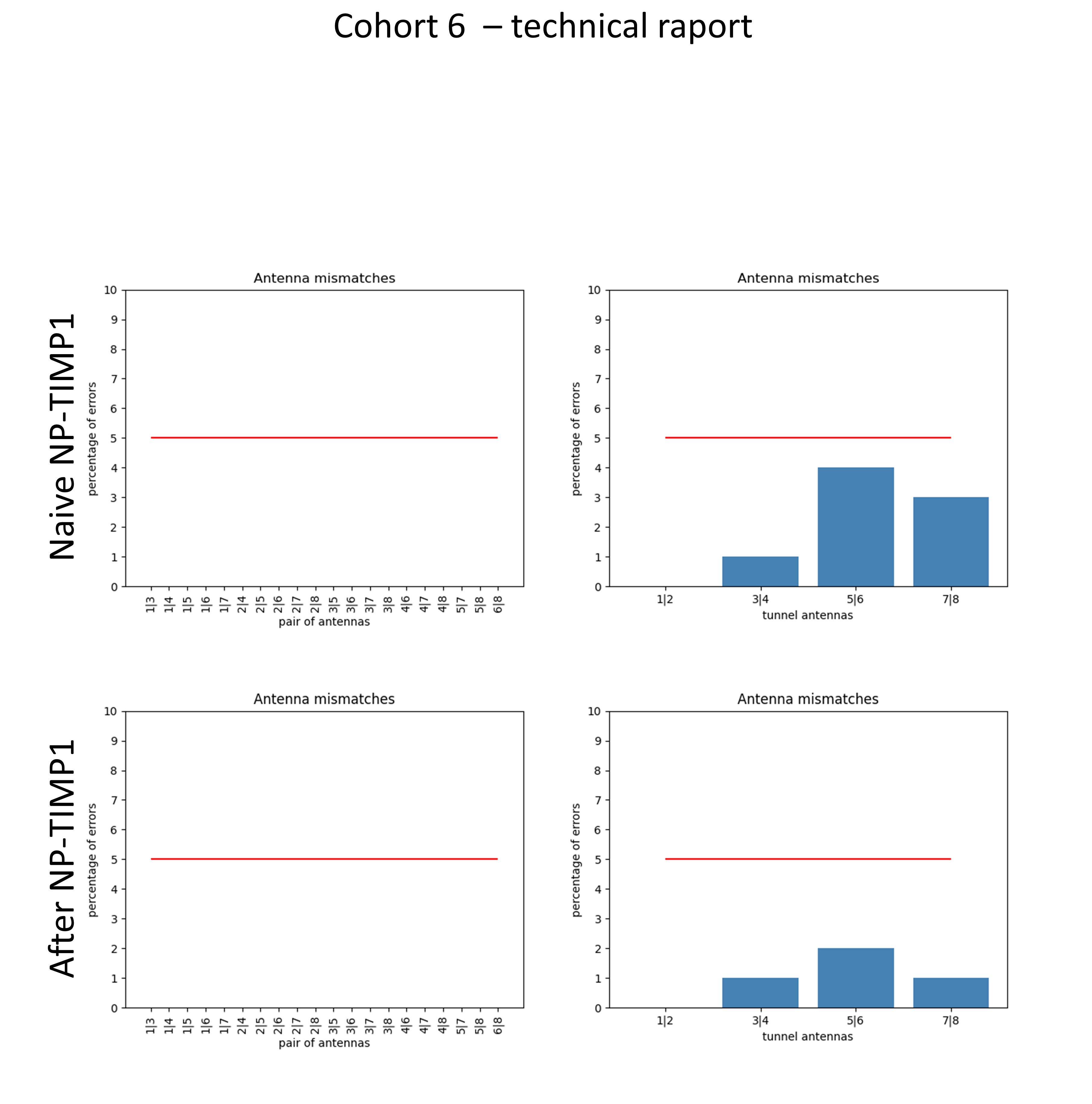

### Fig S12

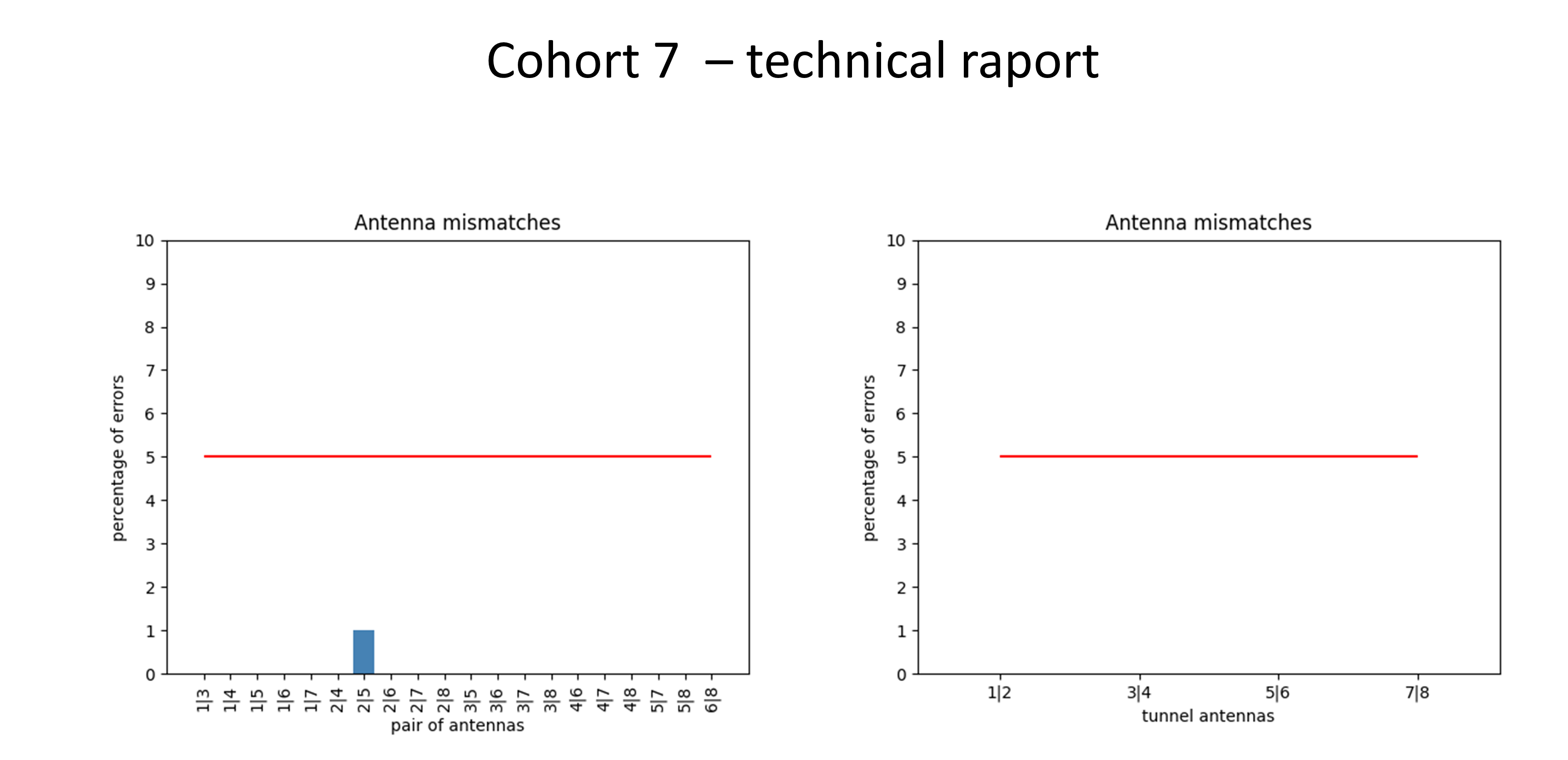
