## Supplementary material for "Social learning about rewards – how information from others helps to adapt to changing environment": Table S1

| Figure | Comparison | Test | Statistic A | Value A | Statistic B | Value B | p-value |
|---|---|---|---|---|---|---|---|
| **Fig.1B** | Approach to social odour - CTRL vs. Approach to social odour- REW n= 10 | Paired t-test | Mean approach to odor | 0,946 +/- 0,07591 | Mean approach to odor | 1,31 +/- 0,1229 | 0,0061 |
| **Fig.1B** | Approach to social odour - REW vs. no change level n=10 | One sample t test | Mean approach to odor | 1,31 +/- 0,1229 | Theoretical mean – no change level | 1 | 0,0324 |
| **Fig.1B** | Approach to social odour - CTRL vs. No change level n=10 | One sample t test | Mean approach to odor | 0,946 +/- 0,07591 | Theoretical mean – no change level | 1 | 0,4951 |
| **Fig.1C** | Persistence in odour seeking - CTRL vs. Persistence in odor seeking - REW n=10 | Paired t-test | Mean persistence | 0,6869 +/- 0,07535 | Mean persistence | 1,079+/- 0,1107 | 0,0294 |
| **Fig.1C** | Persistence in odour seeking - CTRL vs. No change level n=10 | One sample t test | Mean persistence | 0,6869 +/- 0,07535 | Theoretical mean – no change level | 1 | 0,0025 |
| **Fig.1C** | Persistence in odour seeking – REW vs. No change level n=10 | One sample t test | Mean persistence | 1,079+/- 0,1107 | Theoretical mean – no change level | 1 | 0,4930 |
| **Fig.1D** | Followings - CTRL vs. Followings - REW n=10 | Paired t-test | Mean followings events | 0,0507+/- 0.0064 | Mean followings events | 0.1048 +/- 0,0155 | 0,0011 |
| **Fig.2B** | Approach to social odour - REW vs. Approach to social odour - REW-NP-TIMP1 n=11 | Paired t-test | Mean approach to odor | 1,235 +/- 0,07638 | Mean approach to odor | 1,095 +/- 0,1141 | 0,3566 |
| **Fig.2B** | Approach to social odour - REW vs. no change level n=11 | One sample t test | Mean approach to odor | 1,235 +/- 0,07638 | Theoretical mean – no change level | 1 | 0,0117 |
| **Fig.2B** | Approach to social odour - REW-NP-TIMP1 vs. no change level n=11 | One sample t test | Mean approach to odor | 1,095 +/- 0,1141 | Theoretical mean – no change level | 1 | 0,4221 |
| **Fig.2C** | Persistence in odor seeking - REW vs. Persistence in odor seeking - REW-NP-TIMP1 n=11 | Wilcoxon matched-pairs signed rank test | Mean persistence | 1,186 +/- 0,1634 | Mean persistence | 0,7143 +/- 0,0747 | 0,0186 |

| | | | | | | | |
|---|---|---|---|---|---|---|---|
| **Fig.2C** | Persistence in odor seeking - REW vs. no change level n=11 | Wilcoxon Signed Rank Test | Mean persistence | 1,186 +/- 0,1634 | Theoretical median – no change level | 1 | 0,4648 |
| **Fig.2C** | Persistence in odor seeking - REW-NP-TIMP1 vs. no change level n=11 | One sample t test | Mean persistence | 0,7143 +/- 0,0747 | Theoretical median – no change level | 1 | 0,0033 |
| **Fig.2D** | Followings - REW vs. Followings - REW-NP-TIMP1 n=11 | Wilcoxon matched-pairs signed rank test | Mean followings events | 0,0669 +/- 0,0066 | Mean followings events | 0,0329 +/- 0,0025 | 0,0010 |
| **Fig.3B** | Preference for the bottle - CTRL-vehicle n = 9 vs. Preference for the bottle - REW – vehicle n = 10 | Unpaired t-test | Mean relative consumption | 0,7096 +/- 0,07199 | Mean relative consumption | 0,8213 +/- 0,05521 | 0,2297 |
| **Fig.3B** | Preference for the bottle - CTRL-vehicle n = 9 vs. Preference for the bottle - REW – NP-TIMP1 n = 8 | Unpaired t-test | Mean relative consumption | 0,7096 +/- 0,07199 | Mean relative consumption | 0,6162 +/- 0,04403 | 0,3005 |
| **Fig.3B** | Preference for the bottle - REW – vehicle n = 10 vs. Preference for the bottle - REW – NP-TIMP1 n = 8 | Unpaired t-test | Mean relative consumption | 0,8213 +/- 0,05521 | Mean relative consumption | 0,6162 +/- 0,04403 | 0,0130 |
| **Fig.3B** | Preference for the bottle - REW – vehicle n = 10 vs. chance level | One sample t test | Mean relative consumption | 0,8213 +/- 0,05521 | Chance level | 50% | 0,0003 |
| **Fig.3B** | Preference for the bottle - CTRL-vehicle n = 9 vs. chance level | One sample t test | Mean relative consumption | 0,7096 +/- 0,07199 | Chance level | 50% | 0,0196 |
| **Fig.3B** | Preference for the bottle - REW – NP-TIMP1 n = 8 vs. chance level | One sample t test | Mean relative consumption | 0,6162 +/- 0,04403 | Chance level | 50% | 0,0335 |
| **Fig.3C** | Persistence for the bottle - CTRL-vehicle n = 9 vs. Persistence for the bottle - REW – vehicle n = 10 | Unpaired t-test | Mean persistence | 1,012 +/- 0,09684 | Mean persistence | 1,903 +/- 0,3057 | 0,0168 |
| **Fig.3C** | Persistence for the bottle - CTRL-vehicle n = 9 vs. Persistence for the bottle - REW – NP-TIMP1 n = 8 | Unpaired t-test | Mean persistence | 1,012 +/- 0,09684 | Mean persistence | 0,4862 +/- 0,1205 | 0,0037 |

| | | | | | | | |
|---|---|---|---|---|---|---|---|
| **Fig.3C** | Persistence for the bottle - REW – vehicle n = 10 vs. Persistence for the bottle - REW – NP-TIMP1 n = 8 | Unpaired t-test | Mean persistence | 1,903 +/- 0,3057 | Mean persistence | 0,4862 +/- 0,1205 | 0,0012 |
| **Fig.3C** | Persistence for the bottle - REW-vehicle n = 9 vs. no change level | One sample t test | Mean persistence | 1,903 +/- 0,3057 | No change level | 1 | 0,0162 |
| **Fig.3C** | Persistence for the bottle - REW – NP-TIMP1 n = 8 vs. no change level | One sample t test | Mean persistence | 0,4862 +/- 0,1205 | No change level | 1 | 0,0037 |
| **Fig.3C** | Persistence for the bottle - CTRL-vehicle n = 9 vs. no change level | One sample t test | Mean persistence | 1,012 +/- 0,09684 | No change level | 1 | 0,9033 |
| **Fig.3D** | Followings - CTRL-vehicle n = 9 vs. Followings - REW – vehicle n = 10 | Unpaired t-test | Mean followings events | 0,0457 +/- 0,0092 | Mean followings events | 0,0899 +/- 0,0135 | 0,0168 |
| **Fig.3D** | Followings - CTRL-vehicle n = 9 vs. Followings - REW – NP-TIMP1 n = 8 | Unpaired t-test | Mean followings events | 0,0457 +/- 0,0092 | Mean followings events | 0,0213 +/- 0,0069 | 0,0560 |
| **Fig.3D** | Followings - REW – vehicle n = 10 vs. Followings - REW – NP-TIMP1 n = 8 | Unpaired t-test | Mean followings events | 0,0899 +/- 0,0135 | Mean followings events | 0,0213 +/- 0,0069 | 0,0007 |
| **Fig.S1** | Exploratory behavior - CTRL vs. exploratory behavior - REW n = 10 | Paired t-test | Mean exploratory behavior | 635,2 +/- 72,73 | Mean exploratory behavior | 808,1 +/- 109,9 | 0,1134 |
| **Fig.S2** | Exploratory behavior - REW n = 10 vs. exploratory behavior REW – NP-TIMP1 n = 11 | Paired t-test | Mean exploratory behavior | 1086 +/- 94,98 | Mean exploratory behavior | 1076 +/- 117 | 0,8891 |
| **Fig.S3** | Exploratory behavior CTRL-vehicle n = 9 vs. exploratory behavior REW – vehicle n = 10 | Unpaired t-test | Mean exploratory behavior | 892,8 +/- 44,75 | Mean exploratory behavior | 1035 +/- 81,2 | 0,1570 |
| **Fig.S3** | Exploratory behavior - CTRL-vehicle n = 9 vs. exploratory behavior - REW – NP-TIMP1 n = 8 | Unpaired t-test | Mean exploratory behavior | 892,8 +/- 44,75 | Mean exploratory behavior | 545,4 +/- 65,15 | 0,0004 |
| **Fig.S3** | Exploratory behavior REW – vehicle n = 10 vs. exploratory behavior REW – NP-TIMP1 n = 8 | Unpaired t-test | Mean exploratory behavior | 1035 +/- 81,2 | Mean exploratory behavior | 545,4 +/- 65,15 | 0,0003 |
| **Fig.S5** | Correlation followings in Eco-HAB vs. wins in U-tube test | Pearson correlation | Followings | - | Wins | - | 0.0481 |
